# Quantitative mapping of antigen specificity in adaptive immune repertoire embedding spaces

**DOI:** 10.64898/2025.12.09.692930

**Authors:** Marina Frolenkova, Niccolò Cardente, Jahn Zhong, Evgenii Egorov, Giulio Isacchini, Julien Limenitakis, Philipp Fleig, Puneet Rawat, Milena Pavlović, Caterina Sanetti, Jose Gutierrez-Marcos, Geir Kjetil Sandve, Habib Bashour, Maria Francesca Abbate, Victor Greiff

**Affiliations:** Department of Immunology, University of Oslo, Oslo, Norway; Imprint Labs, LLC. New York, NY, USA; Department of Informatics, University of Oslo, Oslo, Norway; Sapienza University of Rome, Rome, Italy; School of Life Sciences, University of Warwick, Coventry, United Kingdom

**Author notes:** Equal contribution (Marina Frolenkova, Niccolò Cardente, Jahn Zhong, Evgenii Egorov). Joint supervision (Habib Bashour, Maria Francesca Abbate, Victor Greiff).

## Abstract

The adaptive immune receptor repertoire (AIRR) encompasses an immense diversity of antibody and T-cell receptor sequences, whose collective organization – how receptors are distributed, clustered, and interrelated across sequence and functional (e.g., antigen-binding) dimensions – remains poorly characterized. Representing AIRRs in continuous representation spaces that capture sequence, biochemical, and structural similarity between receptors may enable comparisons beyond discrete sequence features. Using both one-hot encodings and protein language model (PLM) embeddings, we developed a quantitative framework to map immune receptor organization at global (sequence-set-level) and local (single-sequence-level) scales. Applying the geometry-aware Wasserstein-2 distance, we show that the global structure of the AIRR space can be recovered from as few as ∼10^5^ sequence embeddings, at least 10 orders of magnitude smaller than the theoretical immune receptor diversity. We found that immune receptor sequences annotated with different antigen specificities occupy distinct regions of representation space. To resolve local relationships, we introduce a spatial homogeneity metric that quantifies the extent of functional clustering. We found higher spatial homogeneity in embedding spaces than in sequence space for diverse antigen-specific datasets. Our framework establishes a foundation for quantitative mapping of adaptive immune repertoire organization.

## Introduction

Adaptive immune receptors (AIRs), comprising B-cell receptors (BCRs) and T-cell receptors (TCRs), play a crucial role in the adaptive immune system, mediating antigen recognition and initiating immune responses^1^. Recent advances in sequencing technologies and collaborative efforts within the immunology community have led to an unprecedented accumulation of AIR sequences in public repositories^2–4^. The remarkable diversity of AIRs, generated through V(D)J recombination, and immunoglobulin somatic hypermutations^5^, underpins the specificity and breadth of human adaptive immunity^6–9^. This extreme diversity poses substantial challenges for analyzing and visualizing adaptive immune receptor repertoires (AIRRs), limiting, for example, our ability to characterize how receptors with similar antigen specificity are distributed and organized within the AIRR space.

So far, the majority of analyses have relied on feature-based, repertoire-level approaches that aggregate genotypic sequence characteristics^10,11^ (e.g., sequence similarity, germline gene usage, clonal diversity, or the identification of shared clonotypes – sets of receptors with identical or highly similar CDR3 regions observed across individuals or repertoires). High-dimensional amino acid or sequence-level representations (embeddings) derived from protein language models (PLMs) have proven effective for protein functional annotation, suggesting that their embedding spaces capture information relevant for biological function^12–29^. This has motivated their application to AIR sequence representation^30–32^, mostly using dimensionality reduction approaches in the form of 2D visualizations (PCAs, UMAPs, t-SNE)^31,33,34^. However, to our knowledge, embedding immune repertoires using PLMs has so far mostly reproduced previously known findings, such as (i) AIR sequences cluster by V gene family and sequence similarity^33,35,36^, (ii) BCR sequences cluster by mutational status^37,38^, (iii) AIR sequences cluster by species of origin^35^ and (iv) that antibody sequences are non-uniformly distributed in the sequence space^39,40^. The organization of immune receptors annotated with a similar functional label (e.g., antigen) within the AIRR space remains poorly characterized.

To systematically investigate immune receptor organization across different scales, we define two central concepts: global and local organization. In this work, we use the term “global organization” to refer to sequence-set properties of adaptive immune receptor sequence space. This concept captures how sets of immune receptor sequences are distributed in embedding space beyond clonotypes or germline gene usage. By contrast, we use the term “local organization” to refer to single-sequence-level properties within the AIRR space, capturing how individual receptors with similar functional annotations (e.g., antigen specificity) cluster or colocalize in embedding space. This includes patterns of neighborhood enrichment, functional clustering and spatial proximity between sequences that share functional properties.

While the single-cell field has been fueled by the construction of atlases for novel phenotype identification and estimation of functional distances among cellular subsets^41,42^, such atlases have not yet been built for the adaptive immune receptor domain. Therefore, there is a need to (i) establish strategies that enable systematic comparison of the global organization, diversity and similarity of embedded repertoires across individuals (i.e., collections of sequences from individuals or mixed donors) in a quantitative manner; (ii) investigate whether antigen-specific sequences occupy distinct regions of a human meta-repertoire (composite collection of AIR sequences from multiple donors^43^) distribution within embedding spaces, thereby probing potential biological differences reflected in protein language model representations; and (iii) to quantify the spatial homogeneity of immune receptors with experimentally determined antigen-binding labels within embedding spaces.

In this study, we address these questions by systematically comparing multiple sequence representation strategies (Fig. 1). In addition to interpretable, low-dimensional one-hot encoding (OHE), we evaluate advanced PLMs, including the general ESM2^44^ and the AIR-specific AntiBERTa2-CSSP^45^ (AB2). By applying these complementary approaches to both natural repertoires and antigen-labeled datasets, we show that: (1) BCR repertoires can be efficiently compared using the W₂ distance and that their global organization can be reliably recovered from a limited number of embedded sequences, (2) antigen-specific sequence collections occupy distinct regions within the embedding space, and (3) antibodies sharing identical antigen binding annotation tend to colocalize in close spatial proximity. Our work provides a principled framework for evaluating how embedding spaces reflect functional similarity of adaptive immune receptor repertoires. Of note, in this work, we mainly focused on BCRs, but the methods developed may transfer readily to TCRs.

**Figure 1:**
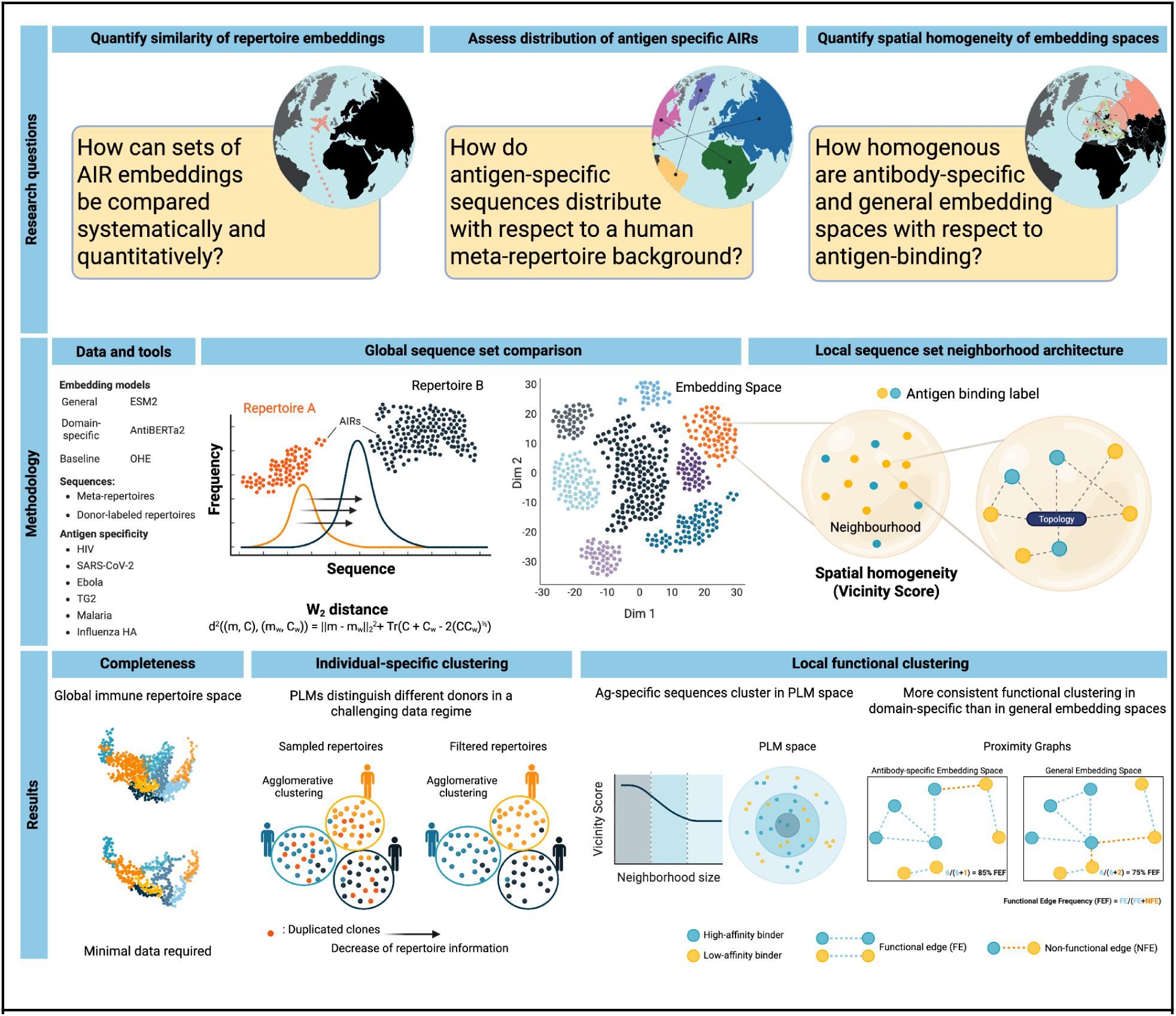
Overview of research questions and computational framework for quantifying global and local features of immune repertoires. We investigated to what extent adaptive immune receptor (AIR) representation spaces reflect biological function at both global (sequence-set-vs-sequence-set) and local (sequence-vs-sequence neighborhood) scales, addressing the current lack of systematic methodologies for such analyses. Using human antibody and T-cell receptor sequences with and without antigen-binding labels from public databases, we generated numerical representations of individual AIR sequences through one-hot encoding (OHE) and protein language models (PLMs), including AntiBERTa2-CSSP (AB2) and ESM2 (see *Methods*). The representations were then aggregated at the repertoire level. Global repertoire organization was assessed using Wasserstein-2 distance, while local functional clustering was evaluated through the vicinity analysis we developed, which examines spatial homogeneity. PLM embeddings reflected meaningful repertoire topology and individual-specific organization, and in several cases revealed antigen-associated clustering, even when data availability was limited. However, in specific contexts, simple sequence metrics like Levenshtein Distance (LD) were observed to perform similarly to, or even outperform, those generated by PLMs.

## Results

### 1. Overview of datasets used in this study

To gain quantitative insight into immune repertoire organization, we compiled a total of ∼22 million human BCR and ∼2 million human TCR sequences from public databases, including iReceptor^3^, Observed TCR Space (OTS)^46^, Immune Epitope Database (IEDB)^47^, and CoV-AbDab^48^. Among the BCRs, ∼18 million are human repertoire sequences labeled by donor ^49^, ∼2 million are unlabeled meta-repertoire sequences^2,3^, ∼450’000 are derived from deep mutational scanning (DMS) assays datasets labeled with binding affinity^50,51^, as well as ∼20’000 antigen-labeled antibody sequences^48^. Of the DMS and antigen-specific datasets, approximately 165’000 and 7’000 BCR sequences, respectively, are paired heavy-light chains, while all other sequences mentioned above correspond to unpaired heavy (BCRs) or beta (TCRs) chains. Comprehensive information about each database is provided in the *Methods* as well as the Supplementary section (Supp. table 1).

## 2. PLM embeddings and Wasserstein-2 distance enable scalable analysis of immune repertoire diversity

### 2.1 Minimal sequence subsets can reconstruct the global structure of immune repertoire representation space

While immune receptor sequence diversity is immense (∼10^9^-10^17^)^49,52–54^, it is fundamentally constrained by the limited genetic variability of its building blocks: the V, D, and J germline genes, and the statistical laws governing V(D)J recombination^40^. Previous work has shown that even small numbers of immune receptor sequences (as few as 1000) suffice to estimate recombination statistics^40^, suggesting that global repertoire organization in sequence space may be recoverable from limited data. Specifically, we asked what the minimum number of embedded sequences required to recover a stable and informative global structure of a meta-repertoire is from a qualitative and quantitative point of view.

To answer this question, we first reasoned that the structure of a meta-repertoire should be reflected in the stability of its embedding distribution. Here, structure refers to the distributional shape of the embeddings, while stability means that this structure remains essentially unchanged as the sample size increases. “Global organization” refers to broad patterns in how sets of antibody sequences occupy the embedding space, shaped by the statistical and biophysical constraints of V(D)J recombination and selection. As more sequences are sampled, the overall distribution of PLM embeddings should gradually converge, assuming that the sampled sequences are representative of the same underlying repertoire distribution. To quantify this convergence, we used the Wasserstein-2 (W₂) distance, an optimal-transport metric that measures the minimal “effort” required to transform one distribution into another^55^. Among existing distance metrics, W₂ has been shown to be particularly well-suited for comparing embedding distributions in generative models, as it captures both geometric and probabilistic differences in a stable and interpretable manner^56^. We measure W₂ between pairs of independently sampled subsets from the same meta-repertoire and expect these values to eventually stabilize, with the stabilization point defining the minimal subset size required to capture global structure.

To assess the smallest sequence subsets sufficient to capture global organization, we randomly selected 300’000 HCDR3 BCR sequences from healthy individuals from the iReceptor database^3^ and generated embeddings using AB2^45^, ESM2^44^ and OHE. These embedded BCR sequences were then divided into batches of increasing size: 10’000-300’000. As a baseline for random structure in PLM space, we generated datasets preserving: (i) amino acid frequency per position (*random_aa*); (ii) total amino acid frequency (*random_shuffled*); (iii) uniform amino acid sampling per position (*random*) and visualized them alongside experimental embeddings using t-SNE^57^, a method widely applied in single-cell transcriptomics to reveal structure in high-dimensional data^58^. t-SNE showed clear organization only in non-randomized BCR data, thereby motivating subsequent quantitative analysis using W₂ (Fig. 2A, Fig. S1).

**Figure 2:**
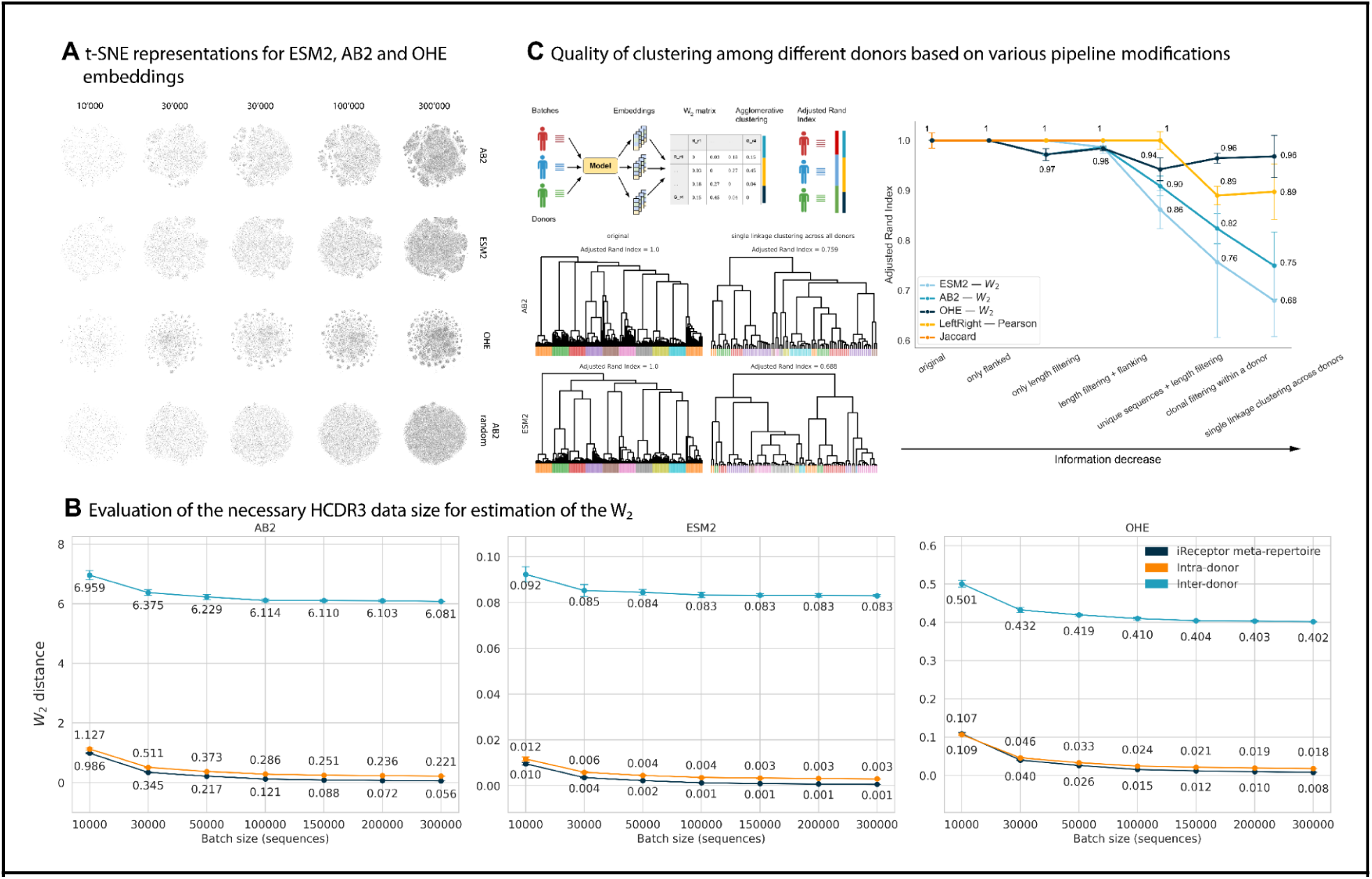
PLMs and W_2_ distance help quantify the organization of embedding space and evaluate its stability. **A** Qualitative visualization of global structure in PLM embedding distributions across increasing sample sizes of the AB2, ESM2, OHE representations for HCDR3 BCR sequences. Each plot shows a t-SNE visualization of the PLM embeddings, with batch sizes noted above and the corresponding model labeled on the right. The global topology of the dataset is discernible starting from 30’000 sequences. **B** Quantitative assessment of global similarity in PLM embedding distributions across increasing sample sizes of the AB2, ESM2, and OHE sequence representations. Each point represents a boxplot of pairwise W₂ distances for each batch size (x-axis) compared to randomly sampled sequences totaling 300’000 from the iReceptor database. Each pairwise distance was calculated ten times to demonstrate that, with a sufficiently large sample, the variance in W_₂_ estimation becomes minimal. **C** Evaluating the quality of clustering replicates (18’000’000 sequences in total) of different donors based on different pipeline modifications. **Upper-left** – scheme of the clusters’ generation pipeline and quality assessment. **Lower-left** – example of the agglomerative clustering applied to all available *Briney et al* replicates. Each tree leaf represents one replicate and it is colored by the respective donors. **Right** – Adjusted Rand Index across all possible model and dataset modifications The ARI for random labels equals 0. **Supplementary figures:** Fig. S1, Fig. S2, Fig. S3, Fig. S4, Fig. S5, Fig. S11, Fig. S12, Fig. S13.

We then embedded 399’990 HCDR3 BCR sequences from the iReceptor database using AB2, ESM2 and OHE, and compared each smaller batch (10’000-300’000 sequences) with the full ∼400’000-sequence set using the W₂ distance. All references to W₂ pairwise distances between batches throughout the text correspond to distances computed between the respective embedding distributions based on their mean and covariance, as detailed in the *Methods* section. For each batch size, we performed 10 independent random samplings to estimate the stability of the embedding distributions (blue line in Fig. 2B). We found that a sample size of 100’000 sequences was sufficient to achieve a near-zero W₂ distance and to accurately estimate the mean and covariance across all embedding types.

To interpret the experiment, it is necessary to determine the expected W₂ distance between samples drawn from the same meta-repertoire distribution. In experimental datasets, technical and biological noise prevent this value from converging to zero, even when samples originate from the same donor. We therefore established baseline W₂ distance values using the Briney et al. dataset^49^, which provides multiple technical replicates per donor. W₂ distances computed between technical replicates (“*intra-donor*”) quantify the minimal divergence expected when two samples represent the same underlying repertoire, while distances between different donors (“*inter-donor*”) capture true biological variation. The “*intra-donor*” baseline thus defines the non-zero convergence level against which pairwise W₂ distances can be interpreted, allowing us to identify the sample size at which W₂ stabilizes to its biologically meaningful lower bound.

For the “*inter-donor*” baseline, we randomly selected one technical replicate per donor, requiring only that it contained a minimum number of sequences (800’000). W₂ distances were then computed across a range of batch sizes (from 10’000 up to 300’000 sequences) between these selected replicates from different individuals. These distances were averaged to yield a representative inter-donor baseline (lightblue line in Fig. 2B). To characterize “*intra-donor*” variability, we similarly selected two technical replicates per donor (also meeting the minimum sequence threshold) and computed W₂ distances between them across the same range of batch sizes. These “*intra-donor*” distances were averaged across all donors (orange line in Fig. 2B).

We found that the W₂ distance between different donors was around 10 times higher than both the distance between technical replicates of the same donor and the distance between different batch sizes from the iReceptor dataset (Fig. 2B). The latter reflects the internal consistency of the same sequence pool when randomly subsampled into batches of varying size, and therefore serves as a baseline for how stable a single embedding distribution remains under repeated sampling. These results suggest that W₂ is suitable for repertoire-level comparisons because it provides an interpretable measure of how far apart two repertoires are in embedding space, integrating many small differences into one coherent signal that corresponds to biological variation.

We performed analogous analysis with ESM2 for TCR data^59^ (Fig. S2) and a DMS dataset of antibodies targeting HER2 (Trastuzumab mutants)^50,51^ (Fig. S3). The results for these two datasets contrast with the findings from the large iReceptor BCR meta-repertoire. For the TCR data, the W₂ distances remained consistently higher across all sample sizes (∼ 3 times higher), suggesting that the global structure of the TCR embedding space is less compact and more challenging to capture reliably with limited sequences. The Trastuzumab DMS dataset showed an even greater deviation, with W₂ distances approximately 10 times higher than the BCR meta-repertoire. This distinct behavior is likely a reflection of the DMS nature, which contains a highly focused set of sequences (Trastuzumab variants) leading to a localized and potentially sparse distribution within the embedding space, further challenging the reliable estimation of distributional distance. These findings highlight that, while the overall distribution of BCR embeddings stabilizes with approximately 100’000 sequences, TCR and specialized datasets, such as the Trastuzumab DMS dataset, may require larger samples or more targeted modeling approaches to achieve similar precision in distributional comparisons.

To summarize, we found that just 100’000 sequences (several orders of magnitude smaller than the estimated diversity of the human BCR repertoire, which is ∼10^9^-10^17,49,52–54)^ are sufficient to describe stable and biologically meaningful distributional properties in PLM embedding space. We also showed that W₂ not only quantifies similarity within and between experimental repertoires, but also effectively distinguishes them from randomly generated controls (Fig. S4).

### 2.2 PLMs reveal donor-specific repertoire structure despite reduced donor-specific information

Having established that PLM embeddings combined with the W₂ metric capture both global repertoire structure and donor-specific signals, we next assessed the robustness (here defined as the ability to retain donor-specific signals) of these representations under conditions that mimic practical limitations of AIRR data, such as low sequencing depth, represented by the absence of identical sequences between the batches or sequences from the same clonal family. Specifically, we asked whether PLM-derived representations of experimental repertoires can distinguish between individuals, even in the more challenging setting in which trivial clonal overlap information is removed. To this end, we analyzed ∼18’000’000 heavy-chain CDR3 sequences from the Briney et al. dataset^49^, which contains multiple replicates from ten donors (18 for each donor), and quantified how well different sequence representations preserve donor-specific signals using the Adjusted Rand Index (ARI)^60^. Briefly, the ARI value reflects how effectively a given encoding-distance combination recovers donor identity, with an ARI of 1 indicating perfect overlap between inferred and true clusters.

We encoded each replicate with five different representations and computed the pairwise distances between all the replicates. In particular, embeddings from (i) ESM2, (ii) AB2 and (iii) one-hot encoding (OHE) were evaluated using the W₂ distance; (iv) sequences encoded with the SONIA left-right encoding model^61^ were evaluated using Pearson correlation; and (v) the Jaccard index, measuring direct sequence overlap. We applied agglomerative clustering on the resulting pairwise distance matrix, constraining the algorithm to the number of original donors (by default, 10) (Fig. 2C upper-left). Visual representations of the resulting clustering structures are shown (Fig. 2C, lower-left, more comprehensive cluster graphs in Fig. S5), where agglomerative trees are colored by the original donor, illustrating the correspondence between inferred clusters and true donor identities.

We then measured how well the inferred clusters matched the original donor labels using the ARI (Fig. 2C right, scheme). To evaluate robustness, we progressively removed sequence-level similarity between replicates, thereby testing how each approach performs as the available information on donor identity becomes increasingly limited (see *Methods*). Across all conditions, the ARI decreased with reduced information, yet OHE and left-right model encoding consistently outperformed PLM-based representations (Fig. 2C). This suggests that, for donor-level repertoire discrimination, simpler sequence-based metrics capture shared clonotypes and overall repertoire similarity better than W₂ distances between sets of PLM embeddings. Nevertheless, PLMs performed above the random baseline (ARI>0.68, consistent randomization of donor labels leads to an ARI = 0.01–0.02), indicating that they retain partial donor-specific information even in the absence of shared clones and the presence of additional data perturbation.

## 3. Qualitative comparison of simulated, natural, and antigen-specific sequences in antibody-specific and general protein embedding spaces

Next, in order to assess how immune receptor sequences are positioned relative to one another in embedding spaces of antibody-specific (AB2) and general (ESM2) PLMs and to determine whether antigen-specific sequences occupy distinct regions of a human meta-repertoire, we compared three classes of sequence collections: (i) a human meta-repertoire (∼2 million sequences) from iReceptor^3^, (ii) simulated repertoires (∼1 million sequences) generated with soNNia^62^, and (iii) seven antigen-specific datasets (HER2^50,51^, HIV^47^, Influenza^47,63^, Malaria^63^, SARS-CoV-2^48^, Ebola^64^, Tg2 (Transglutaminase 2)^65^, total of ∼20’000 sequences, see *Methods* for details). Their sequence diversity was quantified by pairwise Levenshtein distance distributions within and across datasets (Fig. S6).

We computed pairwise W₂ distance matrices using embeddings of the HCDR3 region generated by ESM2 and AB2 (Fig. 3A). Since W₂ is affected by dataset size (Fig. 2B), the diagonal entries were defined as the intrinsic variability of each dataset, obtained by repeatedly splitting repertoires into subsets (see Methods), thus capturing its internal diversity. Accordingly, the W₂ matrices should be interpreted relative to their diagonals: off-diagonal distances greater than diagonal values indicate reliable repertoire differences, whereas comparable or smaller values suggest the difference is indistinguishable from intrinsic noise (see *Methods* for computational details). Antigen-specific datasets (excluding HIV due to very low sample size) were on average ∼6.6 (AB2) and ∼8.7 (ESM2) times more distant from the iReceptor meta-repertoire than the iReceptor meta-repertoire to itself (∼2.00 for AB2; ∼0.02 for ESM2). Simulated repertoires exhibited comparable internal diversity to the iReceptor meta-repertoire, and their distance to the iReceptor meta-repertoire was only ∼1.9-fold their intrinsic W₂ variability, indicating that simulated repertoires are distinguishable from the iReceptor meta-repertoire but show only limited deviation. Notably, AB2 embeddings revealed that Ebola and SARS-CoV-2 datasets were more similar to the meta-repertoire than to the other antigen-specific repertoires (∼3.1 fold and ∼8.9 fold, respectively), whereas this effect was not observed with ESM2 (∼9.2 fold and ∼9.1 fold, respectively). Furthermore, the AB2 distance matrix showed greater variation across datasets, whereas ESM2 produced more uniform distances. After min-max normalization, the variance of AB2 (0.09) was approximately 1.6-fold higher than that of ESM2 (0.05). To summarise, AB2 embeddings yielded larger and more variable W₂ distances between repertoires, providing finer discrimination of antigen-specific differences, whereas ESM2 distances were more compressed and uniform, making repertoires appear more similar overall.

**Figure 3:**
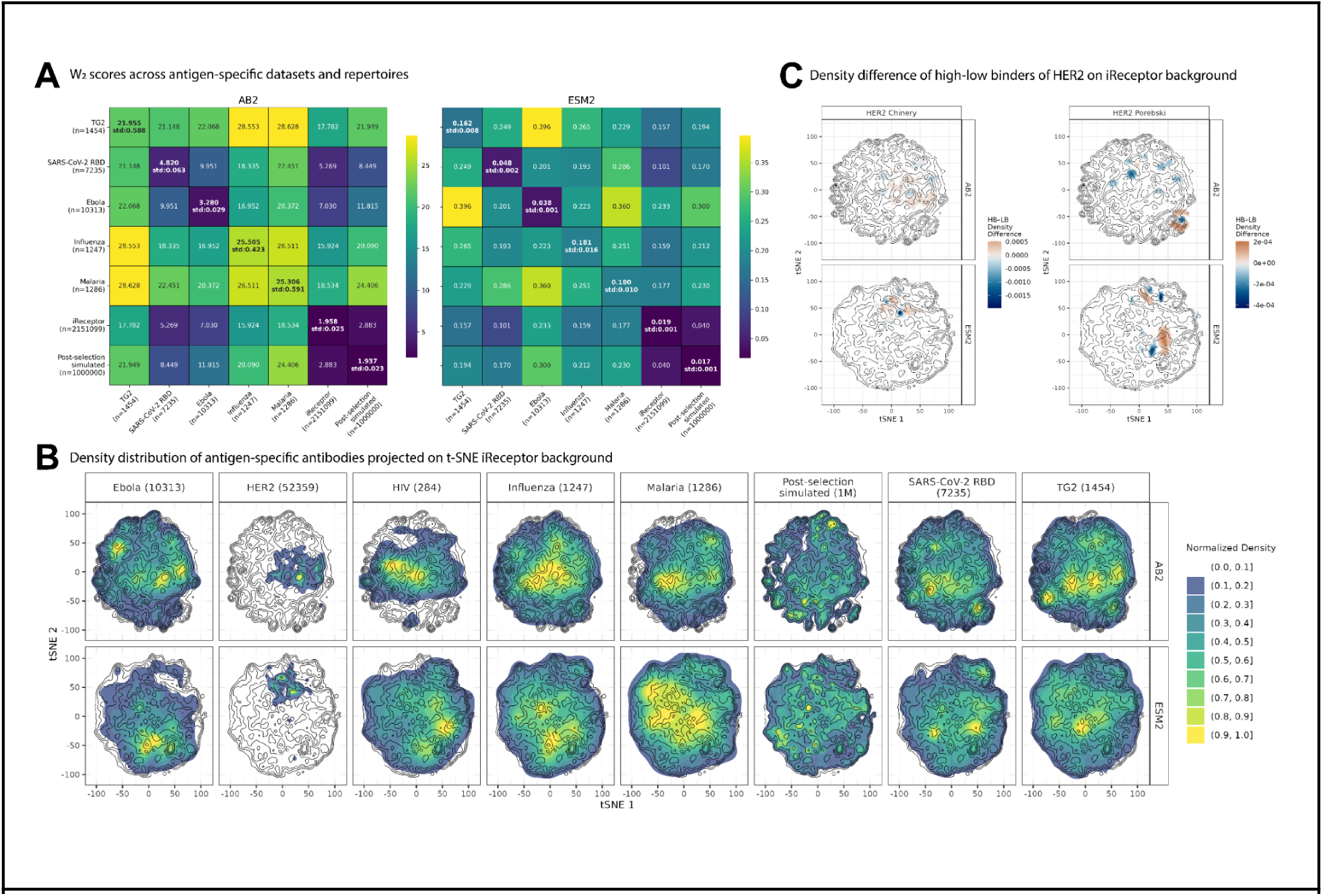
Comparison of sequence-set characteristics across healthy, simulated, and antigen-specific datasets. **(A)** Pairwise Wasserstein-2 (W₂) distances between HCDR3 embeddings (ESM2 and AB2) of a human meta-repertoire (iReceptor), antigen-specific datasets and simulated post-selection repertoires generated from soNNia^62^. For each antigen-specific dataset, the number of sequences is reported in brackets. On the diagonal, the average W₂ distance between two randomly split halves of the datasets is shown. Since this procedure was repeated 10 times with different random splits, the corresponding standard deviation is also reported. **(B)** Two-dimensional projections of antigen-specific datasets using embeddings from two PLMs (ESM2 and AB2). Background contours represent the density of the meta-repertoire, overlaid with the density of antigen-specific datasets. Antigen-specific datasets display distinct density peaks in regions separate from those of the meta-repertoire. To emphasize the regions of highest sequence concentration within each group, densities were scaled by their maximum value (see *Methods*). The colors represent density values scaled from 0 to 1, binned in increments of 0.1. The number of antigen-annotated sequences is shown in brackets. **(C)** Density difference of HER2-Chinery and HER2-Porebski DMS datasets containing high-and low-affinity Trastuzumab variants binding to HER2. To account for differences in HCDR3 length distributions among datasets, each dataset was plotted separately. The 2D projections reveal modest spatially localized increases in estimated probability density for both high-affinity binders (HBs) and low-affinity binders (LBs). HER2-Chinery has 49’721 HBs and 47’328 LBs, whereas HER2-Porebski has 2’638 HBs and 22’152 LBs. Original normalized densities of each binding group are shown in Fig. S7. **Supplementary figures:** Fig. S6, Fig. S7

To visualize the distribution of antigen-specific sequences with respect to the meta-repertoire, we reduced the dimensionality of embeddings to two dimensions by fitting a t-SNE model on the meta-repertoire (Fig. 3B-C). For each antigen-specific dataset, we then computed 2D kernel density estimates (KDEs) in the reduced t-SNE space. In these projections, antigen-specific datasets produced distinct density peaks, mostly unique to each dataset, located in regions separate from the high-density areas of the meta-repertoire background. In contrast, simulated repertoires largely overlapped with the meta-repertoire, suggesting that generative models of V(D)J recombination and selection^40,61,62^ alone accurately characterize the sequence diversity of natural repertoires (Fig. 3B).

To determine whether embeddings capture affinity-related differences within closely related sequences, we focused only on the two deep-mutational scans (DMS) of an antibody (Trastuzumab) targeting HER2^50,51^ (HER2 Chinery, HER2 Porebski) annotated with experimentally determined antigen-binding labels (high- and low-affinity binders denoted as HBs and LBs, respectively, see Methods) and projected them into the same t-SNE space used above (Fig. S7). In both datasets, the two affinity classes occupied broadly overlapping regions and did not form globally distinct clusters, indicating that affinity differences are not reflected at the scale of the entire embedding space. To investigate a more subtle structure, we compared their normalized 2D KDEs by subtraction (Fig. 3C, see Methods). This revealed only small, localized density shifts (more noticeable in the Porebski dataset and minimal in the Chinery dataset), indicating that any affinity-dependent structure is limited to small subregions and lacks a consistent spatial pattern relative to the meta-repertoire background. Together, these results show that while antigen-specific datasets separate in the embedding space, affinity differences within a DMS dataset yield only weak, dataset-dependent local enrichments rather than global separability.

To quantify spatial organization more rigorously, we performed spatial homogeneity and topology analyses directly in the full embedding space rather than in the t-SNE-reduced space (next section).

## 4. PLM embeddings reveal local antigen-specific organization reflected by the Vicinity Score

While the 2D t-SNE projections of PLM embeddings (Fig. 3C) provided a helpful overview of the separation of high and low-affinity binders in the embedding space, their inherent limitations led us to pursue a more rigorous, quantitative analysis. Our focus now is to compare the extent to which antibody-specific and general PLM representation spaces, as well as the baseline sequence space, capture functional (i.e., antigen-binding) relationships among AIRs.

Prior to the advent of PLMs, functional similarities of AIRs were commonly approximated using edit distance metrics such as Levenshtein distance (LD) and its variants^66–68^. While useful for antigen-binding prediction^69–72^, LD in sequence space is potentially ill-suited for topological or functional comparisons: its discrete scoring treats all amino acid substitutions equally, regardless of physicochemical similarity, obscuring structural and functional consequences compared to more sophisticated distance measures, such as TCRdist^73^. Resolution further degrades as neighborhood size grows, since many sequences converge to the same distance despite substantial biological differences. In contrast, PLMs embed sequences into continuous similarity spaces where distances vary smoothly^74^, can be thresholded with high precision, and allow efficient computation of proximity graphs^75^ via vectorized metrics such as Euclidean or cosine distance. Whereas LD complexity scales quadratically with sequence length^76^, the complexity of Euclidean- and cosine distance between PLM embeddings scales linearly with length. Thus, PLMs may offer both physicochemical interpretability and computational efficiency, potentially enabling finer discrimination of functional similarity than discrete sequence space.

Given the potential capacity of PLMs to capture structural and functional characteristics of proteins^44,45^, we asked whether AIRs with similar function (in this case target binding) colocalize in PLM embedding space (spatial homogeneity) and whether distances in embedding space better reflect functional differences than those in sequence space^12,77^. We tested our hypothesis across two spatial scales: (i) assessing spatial homogeneity of antigen-binding within local neighborhoods in both sequence and embedding spaces; and (ii) constructing graph-based representations to evaluate the functional topology of these spaces, specifically comparing the fraction of shared antigen-binding labels between adjacent sequence embeddings.

To quantify local antigen-specific clustering in PLM embedding space (i), we developed the *Vicinity Score*, a measure of spatial homogeneity that captures the tendency for nearby sequences in embedding space to share functional labels such as antigen specificity. The Vicinity Score is defined as the median precision across sequences, where precision is here the fraction of sequences sharing the same label (e.g., target antigen) within a given cosine distance radius of an embedded AIR. Thus, both precision and the Vicinity Score are defined as functions of the radius. To prevent biases due to uneven neighborhood densities, we include a resampling step that weights precision values inversely by local sequence density (Fig. 4A, see *Methods*). The Vicinity Score measure can be interpreted as asking whether a hypersphere drawn in embedding space around a given antibody is enriched for functionally similar antibodies. In this study, the functional label refers to high-affinity binders (HB); all Vicinity Scores reported in the main figures are therefore computed on the HB class.

**Figure 4:**
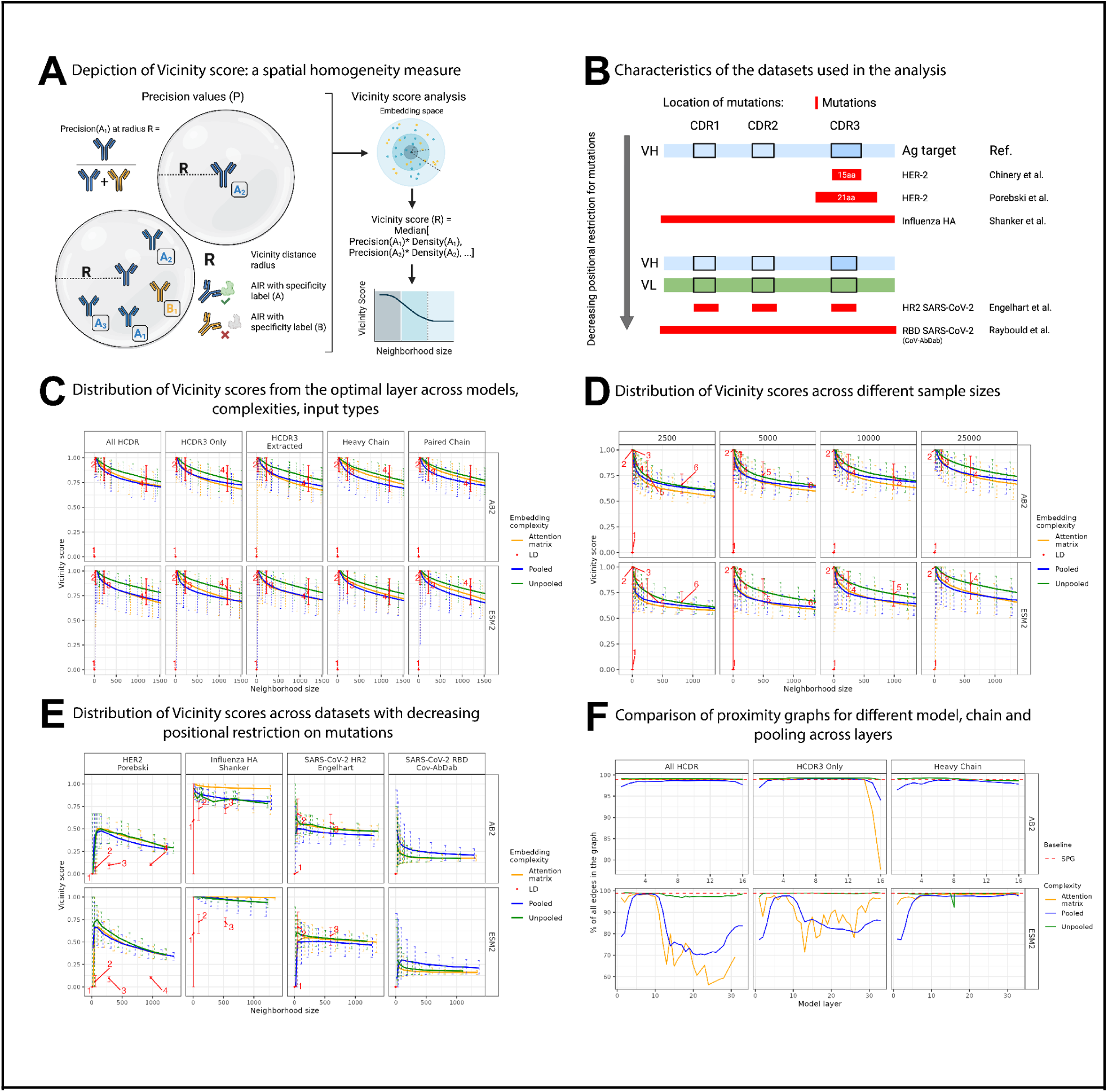
Unsupervised representations are able to spatially segregate similar binder sequences in a continuous space. (**A**) Graphical representation of the Vicinity Score: For each AIR, represented by a point in the space, we compute the precision (defined as the fraction of points with the same HB label as the initial point, within a specified cosine distance radius R). For each radius R, we take the median of the Precision values sampled by their Local Density (see *Methods*). (**B**) Characteristics of the datasets used for vicinity (A-E) and graph analysis (F), ordered from positions with the highest restriction on mutational variation (mutations in HCDR3 only) to those with little or no restriction. The target antigens of the selected antibodies are HER2^50,51^, influenza hemagglutinin (HA-H1)^78^, SARS-CoV-2 HR2 region^79^, and SARS-CoV-2 RBD (collected from Covabdab^48^). The red bar indicates the antibody region where mutations are located (see *Methods*). “Ref” corresponds to the first author of the dataset’s original publication. (**C-E**) Distribution of Vicinity Scores as a function of the neighbourhood size. The dataset was embedded using combinations of variables, including embedding complexity, sequence context, and model (see *Methods*). For each combination of model, embedding, and sequence context, only the optimal layer *l\** as determined by the highest *mc* is displayed. As a baseline, Vicinity Scores were also computed with Levenshtein distance. Numbers in red indicate the LD (in amino acid) threshold. The line colors indicate the embedding complexity used for Vicinity Score calculation. (**C**) HER2 Chinery dataset of 97’049 total sequences: 49’721 high-affinity binders and 47’328 low-affinity binders. (**D**) HER2 Chinery dataset subsampled in 2500, 5000, 1000, 25000 sequences per binding label. The data represented is from the “HCDR3 only” sequence context variants. (**E**) Data are shown from multiple different datasets (DMS and real-world data from CoV-AbDab) with a decreasing level of positional restriction on mutations. See *Methods* for a detailed description of each dataset. (**F**) Embedding proximity graphs (EPGs) contain a higher proportion of edges between sequences with the same binding label compared to a sequence proximity graph (SPG). EPGs are constructed from PLM embeddings from (C) and compared to SPG constructed from sequences at LD = 1. Each node in EPG represents a sequence embedding. Nodes with a cosine distance below a threshold are connected by an edge. The threshold is defined as the average cosine distance between embeddings of sequences with LD = 1. In each panel, the proportion of same-label edges in the SPG is indicated by a black dotted line. **Supplementary figures:** Fig. S8, Fig. S9, Fig. S10, Fig. S14, Fig. S15.

We computed the Vicinity Scores for multiple datasets with varying characteristics^48,50,51,78,79^, ranging from deep mutational scanning (DMS) datasets with mutations restricted to HCDR3, to antibody datasets mutated across full heavy chains or paired heavy and light chains. These datasets differ in positional restriction of mutations, sequence length, and complexity (Fig. 4B, see *Methods*). For each dataset, we computed the Vicinity Score across five context window variations corresponding to the columns in Fig. 4B: (1) heavy chain CDR1–3 concatenated (All HCDRs), (2) HCDR3 only, (3) full heavy chain with only the CDR3 tokens extracted (HCDR3 Extracted), (4) full heavy chain (Heavy Chain), and (5) full paired chain (Paired Chain). For each variation, we evaluated embeddings from two PLMs (ESM2 and AB2; rows in Fig. 4B) under multiple embedding complexities—mean-pooled (hereafter, pooled), unpooled, attention matrices—alongside the Levenshtein distance baseline, producing one Vicinity Score curve per configuration. The Vicinity Score was applied across 15 progressively wider radii, allowing a comprehensive assessment of functional clustering in diverse embedding spaces. As absolute cosine distances are not directly comparable across embedding spaces, Vicinity Scores were computed across multiple neighborhood sizes (see Methods).

Although final-layer embeddings are usually chosen for downstream analysis^17,80,81^, our layer-by-layer evaluation showed that this is not always optimal for capturing the colocalization of functionally similar antibodies. We summarized each curve by its mean Vicinity Score across radii (*m_c_*), and for each model, selected the layer with the highest *m_c_*. We observed that layers exhibiting the highest *m*_c_ (Fig. S8), are predominantly (80%) distributed in the first half of the layers (1–16 for ESM2, and 1–8 for AB2). All subsequent analyses were conducted at this optimal layer depth *l\** , with *m_c_* being used to summarize curves and average effect sizes to quantify differences (see *Methods*).

We begin by illustrating the Vicinity Score (Fig. 4C) on a subsample of the 349’893 Trastuzumab DMS variants (targeting HER2) dataset^50^, comprising unique 49’721 high-affinity (HBs) and 47’328 low-affinity binders (LBs). We compared pairwise Levenshtein distances and showed that the distance distributions between pairs with the same label (HB-HB or LB-LB) substantially overlap with the distribution of HB-LB distances (Fig. S9), suggesting that sequences of the two binding classes are not trivially separated by Levenshtein distance. We focused all our subsequent analysis only on the high-affinity binder group.

Within the HB class, HB Trastuzumab variants are predominantly neighbored by other HBs. This is reflected in the high Vicinity Scores we observed for small neighborhood sizes (with Vicinity scores values ranging from ∼1.0 to ∼0.85, across the first 250 neighbors), indicating antigen-associated colocalization in embedding space. At the same time, the large variability (IQR typically between ∼0.034 and ∼0.4, with some outliers reaching ∼1) highlights that HB label homogeneity in local neighborhoods can vary substantially across the embedding space. When compared to the LD metric, PLM embeddings showed comparable performance: unpooled embeddings were nearly indistinguishable from LD (average effect sizes ∼0.02–0.12), attention matrices were moderately comparable (∼0.07–0.08 for AB2 embeddings of heavy and paired chains, and ∼0.21–0.39 in other cases), while pooled embeddings showed the largest differences (∼0.25–0.37). All effect sizes can be found in Supp. table 2. Our analysis indicates that embedding complexity has a stronger influence on spatial homogeneity than the amount of sequence context provided. In other words, pooling of embeddings impacted the Vicinity Score more than extending the input context window from HCDR3 alone to the full heavy chain or even paired chains, as long as mutations remain confined to the HCDR3. Further, even in the best case, PLM embeddings matched or lagged behind the LD baseline’s Vicinity Score.

To assess robustness of the Vicinity Score to dataset size, we repeated the analysis on the Trastuzumab variants dataset using the (2) HCDR3 only input, subsampling high and low-affinity binder classes to 2’500, 5’000, 10’000, and 25’000 sequences (Fig. 4D). Vicinity Scores for both PLM-based and LD-based methods decreased with smaller samples, but the performance gap between the best PLM method and LD remained stable, confirming the same trend of Fig. 4B. In LD space, smaller samples shifted the effective neighborhood range toward higher LD values due to the scarcity of closely related variants—a statistical artifact of sparser sampling rather than a biological effect. Across all sizes, unpooled embeddings maintained higher Vicinity Score distributions than attention matrices or pooled embeddings, yet remained comparable to the LD baseline.

We next evaluated the generalizability of our findings by expanding the analysis to different antigen-binding labeled datasets, each characterized by a decreasing positional restriction for mutations (Fig. 4E, *Methods*): Trastuzumab DMS variants targeting HER2 from Porebski et al.^51^, mutated across 21 aa of HCDR3; CR119 Influenza bnAb anti-HA variants from Shanker et al.^78^ mutated across the full heavy chain; SARS-CoV-2 (directed towards HR2-spike region) antibody variants from Alphaseq^79^, generated by mutating CDRs of either heavy or light chains; and SARS-CoV-2 antibodies from CoV-AbDab^48^ only directed towards RBD, paired with background non-binders from public antibody databases^2^. Overall, Vicinity Scores were lower than in the first Trastuzumab DMS dataset^50^ (*m_c_* ∼ 0.75 for HER2-Porebski^51^, 0.55 for SARS-CoV-2 HR2, 0.45 for SARS-CoV-2 RBD) but still indicated spatial homogeneity of binder labels. Notably, the Influenza HA dataset^78^ yielded exceptionally high scores with ESM2 attention matrices (*m_c_*= 1), while LD space in some datasets—especially SARS-CoV-2 RBD from CoV-AbDab—did not display any spatial homogeneity (*m_c_* ∼ 0). In contrast, the SARS-CoV-2 HR2 Alphaseq dataset favored LD over PLM embeddings (effect size = 0.27 -0.40), likely due to its library design with minimal point mutations from each parental sequence (see *Methods*). Across these datasets, our earlier finding—that final transformer layers are not consistently optimal for the spatial homogeneity task we have been evaluating—remained valid (Fig. S10).

Across our experimental setups (Fig. 4B-D), we observed that the unpooled embeddings yielded generally higher Vicinity Scores than the pooled embeddings, highlighting the importance of preserving residue-level information. In fact, mean pooling appears to discard critical positional information necessary for capturing functional specificity, particularly relevant in the antibody field, where a single amino acid change at a localized position may alter binding reactivity towards the antigen ^50,82^.

### 4.1 Topology of antibody-specific embedding proximity graphs captures functional adjacency more consistently than general embedding proximity graphs

Having observed that the association between embedding distance and function was higher in more localized regions of embedding space (Fig. 4C-E), we set out to analyze the topological relationship between *immediately* neighboring (embeddings of) sequences (Fig. 4F). To that end, we began by constructing embedding proximity graphs (EPGs) and a baseline sequence proximity graph (SPG) for Trastuzumab DMS variants, as described in Chinery et al.^50^ (see Methods). One EPG was constructed for each context window variation (“All HCDRs”, “HCDR3 only”, “Heavy chain”), output type (pooled/unpooled embeddings, attention matrix), embedding model and layer (AB2: 16 layers, ESM2: 33 layers). Briefly, sequences are represented as nodes in a graph, and unweighted edges connect sequences or embeddings that are adjacent to each other. Each node is labeled as a high- or low-affinity binder (HB or LB). Sequences in the SPG are considered adjacent if they are exactly 1 LD apart. Embeddings in EPGs are defined as adjacent if their cosine distance is below the average distance of embeddings whose sequences are 1 LD apart. Edges between nodes of the same label (HB/HB or LB/LB) are “functional edges” (FEs), while edges connecting HB and LB labelled nodes are “non-functional edges”. For the SPG and each EPG, we calculate the functional edge frequency (FEF), defined as the percentage of functional edges among all edges in the graph.

As shown in Fig. 4F, the baseline SPG already had a near-perfect FEF of 98.89%. The FEF was comparably high in EPGs constructed from AB2 embeddings, regardless of context window variation, pooling, model layer or domain (∼96-99%; see Fig. 4F), with the only exception of “HCDR3 only” representations from its last layer (Layer 16; FEFs unpooled: 98.86%, pooled: 94.01%, attention matrices: 77.66%). Although general EPGs from unpooled ESM2 embeddings also match the FEF of AB2 and SPG baseline across all 33 model layers and context window variations, EPGs from pooled ESM2 embeddings exhibit low FEFs of ∼80% that increase to a plateau of ∼95% from layer 5 onwards. While the FEF remained high until the last layer (layer 33) in EPGs from pooled “Heavy chain” embeddings of ESM2, FEFs of pooled “All HCDRs”-and “HCDR3 only”-EPGs are only high in intermediate layers (layers ∼5-10). Our findings indicate that the antibody-specific model (AB2) represents antibodies in a way that preserves their functional relationships across embedding layers, even when supplying alternative context window variations. Further, although EPGs from the general PLM (ESM2) are able to match antibody-specific EPGs in some cases, FEF in general EPGs is much more sensitive to pooling, alternative context window variations and choice of embedding layer.

Collectively, our vicinity and topological analyses indicate that, although Levenshtein distance (LDs) is generally ill-suited for capturing functional relationships, it performs well in cases where datasets have low sequence diversity (Fig. 4C-D,F), likely reflecting effects from DMS dataset construction methods that inherently follow LD structure by sampling sequences that are similar to known binders. In these instances, protein language models (PLMs) tend to match or fall short of the LD baseline. Conversely, in high-sequence-diversity settings where LD fails to capture functional similarity, PLMs demonstrate a superior ability to colocalize antibody sequences based on their binding labels.

## Discussion

### 1. Rationale for using the Wasserstein-2 distance to compare AIRR embedding distributions

When immune receptor datasets are represented as continuous distributions in a high-dimensional representation space, comparing immune repertoires (or AIR sequence sets) reduces to comparing their overall distributional structure rather than individual sequences. The W₂ distance is well-suited for this comparison task, as it quantifies global shifts between distributions and has been widely validated in analogous settings, such as the Fréchet Inception Distance (FID)^83^, Fréchet Audio Distance (FAD)^84^, and Fréchet Video Distance (FVD)^85^.

To date, no single metric is consistently used across AIRR studies to compare BCR or TCR embedding distributions. Motivated by this objective, we investigated and, upon validation, adopted the Wasserstein-2 distance (W₂) as a metric for quantifying differences between multivariate embedding distributions. Similar optimal transport (OT) based frameworks have recently been applied to TCR repertoire analysis^86^. Olson et al.^86^ first encoded TCRs using the TCRdist^73^ framework, which measures receptor pairwise distances based on their CDR loops, with mismatch scores derived from BLOSUM62 plus an extra weighting for CDR3 regions. The resulting distance matrix, rather than high-dimensional embeddings from PLMs, was used as input to OT, enabling repertoire-level comparisons despite OT’s computational cost, demonstrating that OT metrics can capture biologically meaningful differences between repertoires without requiring explicit modeling assumptions. While TCRs are structurally and functionally related to B-cell receptors (BCRs), they lack the somatic hypermutation process that drives variability in BCR repertoires, making BCRs uniquely challenging for embedding-based similarity assessments. Previous repertoire comparison frameworks such as immuneREF^10^ quantify repertoire similarity by integrating predefined immunological features. In contrast, our Wasserstein-2 approach operates directly in the continuous embedding space derived from protein language models, allowing unbiased quantification of repertoire-level distances without requiring explicit feature selection.

While most applications of the Wasserstein-2 metric have focused on generative modeling in images or molecules, recent studies have begun applying similar distributional analyses to protein language model embeddings^87^. Rissom et al.^87^ introduced a general framework for the quantitative comparison of embedding spaces in protein language models, demonstrating that global and local distance statistics can capture biologically meaningful variation across models and datasets. Our findings support the interpretation of metrics such as the W₂ distance for repertoire-level comparison as informative measures of how well embedding distributions preserve biological structure.

### 2. Vicinity score quantifies spatial homogeneity in high-dimensional spaces

Although protein language models (PLMs) have shown promise for learning high-dimensional embeddings of proteins^21,45,88^, their applications in biotechnology are still emerging, especially when compared to the scale of commercial large language models with vastly more parameters and number of tokens in their training datasets (e.g., ChatGPT^89^, Gemini^90^, Claude^91^)^92–94^. Nevertheless, PLM embeddings have proven effective for predicting protein structure and functions^15,44,78^, suggesting that spatial proximity in embedding space may serve as a proxy for functional similarity. However, these analyses were conducted on general protein datasets spanning multiple species, where sequence homology often reflects functional similarity. AIRs, by contrast, represent a more complex case, as their function (binding) is defined with respect to a given antigen and their sequence variability is mostly restricted to the relatively short CDRs where even a single amino acid change can drastically alter binding and therefore function, making homology a less reliable predictor for antigen-binding specificity.

Independently and parallel to our research, recent studies^16,95,96^ attempted also to quantify the spatial homogeneity of PLM embedding spaces in relation to functional annotation labels. These methods have in common that they define the neighbors of a given sequence (k-nearest neighbor approach, KNN) and then examine the proportion of the sequence’s neighbors with the same functional label as itself. Most current methods for analyzing distances in embedding spaces rely on KNN^16,87,97^. While effective in a single and well-defined space, this approach has two key limitations. First, embedding spaces are sparse and non-linear, such that the distance to the *k*^th^ neighbor depends heavily on local density, reducing the generalizability of conclusions^98^. Second, the density of embeddings varies across models and layers, making the definition of a fixed neighborhood imprecise across spaces. To address these issues, the Vicinity scoring method that we developed is computed with radius-based neighborhoods defined by a common rule across models and layers. Additionally, we implemented a *density-weighted resampling* step to better account for sparsely distributed sequences, ensuring that underrepresented regions contribute proportionally to the analysis. While these precautions improve the robustness of embedding-space comparisons, substantial work remains to fully address confounding factors arising from variations in density and differences across embedding spaces.

Our analyses demonstrate that PLM embedding spaces can spatially cluster sequences with identical binding labels, achieving Vicinity Scores comparable to or superior to those of the Levenshtein distance (LD). Across multiple datasets (Fig. 4C-E), unpooled embeddings generally yielded higher Vicinity Scores than pooled representations, highlighting the importance of retaining residue-level or positional information. Mean pooling appears to dilute critical positional signals, which is particularly relevant in antibodies, where single-residue substitutions can determine antigen binding. Exceptions were observed in datasets with non-uniform amino acid lengths (Fig. 4E). We hypothesize that sequence-length variability, requiring padding in unpooled embeddings and raw attention matrices, may introduce alignment noise in cosine-distance calculations and confound spatial comparisons. Addressing variable sequence lengths with more sophisticated approaches will likely be necessary to fully exploit residue-level embeddings.

Previous studies have shown that the final layers of PLMs consistently underperform on downstream tasks, with intermediate layers typically yielding the best results^99–101^. We observed a similar trend; however, in our case, the early layers performed best. Since our analysis focuses on the spatial organization of the embedding space, this finding aligns with the detokenization hypothesis, which suggests that early layers primarily transform raw tokens into a structured embedding space.

Finally, results from the HER2 Chinery experimental data (Fig. 4C) suggest that incorporating sequence context from non-mutated regions is less informative than retaining single-residue information from mutated regions. In AB2, embeddings appear highly redundant, as strong Vicinity and topology scores were achieved across layers even after pooling. However, alternative explanations are also possible. For example, AB2 may concentrate functionally relevant variation in fewer dimensions, making its representations more robust to pooling. In contrast, redundancy in ESM2 is lower: unpooled embeddings yield high Vicinity Scores, whereas pooling restricted to CDRs, particularly in early layers, results in poorer performance (Fig. 4E, Fig. S8). Overall, our findings indicate that reducing residue- and position-specific information through pooling reduces spatial homogeneity of PLM embedding spaces more than reduced sequence context, with AB2 showing greater embedding redundancy than ESM2.

## 3. Limitations of 2D projections, PLM-based representation, W₂ estimation and sequence datasets

### 3.1 Reliability of 2D projections and PLM-based repertoire signatures

T-SNE and UMAP projections provide intuitive overviews of repertoire organization, but they inevitably distort high-dimensional structure^102,103^. We observed that antigen-specific sequences form clear, separate peaks in embedding space, which could serve as reference points for comparing antibodies against the same target. But it is worth noticing that apparent separation or clustering in two-dimensional maps may also reflect compression artifacts or amplification of technical biases. Direct analyses in high-dimensional embedding space, such as using W2 distance (Fig. 2, Fig. 3) or Vicinity Scoring (Fig. 4), therefore, remain essential for rigorous interpretation of repertoire similarity.

Our analyses reveal that PLM-derived representations of immune repertoires are shaped by multiple sources of bias, ranging from intrinsic sequence properties and data sampling^104^ to choice of similarity metrics and embedding variables.

A key step in evaluating the PLM embeddings and their limitations is determining what biological information PLMs encode. For example, PLM embeddings may capture functional aspects of immune repertoires that correspond to known biological properties, such as donor immunogenetic features, repertoire diversity, and antigen specificity. Our results demonstrate that PLMs can reliably capture donor-level variation (Fig. 2C) and distinguish natural repertoires from randomized controls (Fig. S4), indicating that they encode meaningful biological signals. Even after stringent filtering (removing identical sequences, clonal families, and fixed-length subsets) donor identity remains detectable in PLM embeddings (Fig. 2C, Fig. S5). However, the quality of replicate clustering is limited, suggesting that PLMs capture a weaker repertoire-level signature compared to simpler encodings such as OHE, SONIA left-right encoding model-based features^10,61^, or Jaccard-style overlap metrics. We hypothesize that these simpler methods concentrate the signal on compositional or positional statistics that remain after filtering, whereas mean pooling and W₂ computation in PLMs may smooth fine-grained contextual cues, potentially reducing discriminative power.

It is important to note that the biological signal captured by embedding representations depends strongly on technical choices such as preprocessing steps, sequence window context^105^, layer selection, and pooling strategy^106^, all of which can substantially alter how repertoire structure is expressed in the representation space (Fig. 4). These factors must be carefully considered to ensure that the biological signals in embedding space are correctly identified and not confounded by technical choices. For tasks such as donor monitoring or stratification, simpler encodings may therefore remain more robust. Beyond amino-acid level embeddings, recent work has shown that models explicitly incorporating nucleotide context and selective pressures can capture evolutionary and mutational patterns in BCRs that are not fully represented in standard PLM embedding spaces,^23,29^, pointing toward a new generation of repertoire-aware sequence models.

### 3.2 Embedding distance metric

There are various metrics that measure the similarity or distance between two embeddings. We adopted cosine distance as the primary metric for defining neighborhood relationships (Fig. 4A-F), as it is widely used in analyses of protein language model (PLM) embeddings. Its scale invariance ensures robustness across layers and facilitates comparability with prior work^14,15,107^, and its computational efficiency makes it suitable for large-scale repertoire analyses.

However, cosine distance only measures the angular difference between vectors and does not account for the magnitude, which can be limiting if vector norms carry biologically meaningful information (e.g., embedding confidence or contextual strength). In high-dimensional spaces, cosine distances can also concentrate, reducing their ability to capture fine-grained similarities^108,109^. Alternative metrics such as Euclidean distance, Manhattan distance, or embedding-based alignments (EBA) could better capture functional relationships^14,110–113^ and may therefore yield different embedding-space organization and proximity graph topologies. Notably, EBA inherently accommodates variable sequence lengths and detects homologues sharing less than 20% sequence identity, whereas traditional metrics can be distorted by padding tokens. Yet, EBA is not as scalable as cosine distance due to its computational complexity and, to our knowledge, has not been applied to functional antibody repertoires, where sequence homology is a poor proxy for antigen-binding similarity. Future work should therefore explore combinations of distance metrics and alignment-aware approaches to better characterize antibody functional landscapes in embedding space.

### 3.3 Dataset size and W₂ estimation

Dataset size constrains the robustness of downstream statistical analyses. In fact, reliable computation of global metrics such as the Wasserstein-2 distance (W₂) depends on stable estimates of means and covariances. When the sample size is small relative to the embedding dimensionality, the covariance matrices become noisy and W₂ values are unstable. In our analyses, convergence was achieved only for datasets exceeding ∼10^5^ sequences, whereas smaller subsets produced inconsistent values (Fig. 2, Fig. S2, Fig. S3), a limitation noted in other high-dimensional domains applying Wasserstein-based metrics^55^. These effects reflect statistical ill-conditioning rather than biological variability. For limited datasets, W₂ should thus be interpreted cautiously and, where possible, complemented by regularized or resampling-based approaches.

While efficient and interpretable, W₂ estimation is also sensitive to the fraction of shared or highly similar sequences between datasets. As shown in our analyses, W₂ decreases non-linearly as more sequences are shared between two repertoires (Fig. S11). This nonlinearity indicates that W₂ partially reflects data redundancy, and its interpretation requires careful consideration of clonality and public clones. Without these adjustments, W₂ may overestimate similarity in repertoires rich in shared sequences, while overestimating dissimilarity in those with few.

### 3.4 Intrinsic sequence properties

First, the length of the HCDR3 sequence strongly influences PLM organization: sequences of the same length cluster together in embedding space, an effect that persists even in random controls matched for length distributions. As a result, local neighborhood structure can be dominated by length rather than biological similarity, making repertoires appear more separated or homogeneous in low-dimensional projections than they truly are (Fig. S12). This motivates strict length control when comparing repertoires (e.g., fixed-length subsets or length-stratified analyses) and, where possible, the inclusion of flanking residues (C/W amino acids) to standardize boundaries.

A second intrinsic bias arises from germline gene composition. Even when embeddings are restricted to HCDR3 regions, receptors derived from the same J or D gene families form distinguishable clusters (Fig. S13), consistent with prior reports that PLM-derived embeddings retain V(D)J gene-dependent structure^13^. This germline-driven localization suggests that PLMs encode sequence origins as well as functional properties, potentially exaggerating or masking biologically relevant similarities—e.g., repertoires differing in gene usage may appear distant despite comparable functional diversity.

### 3.5 Biases in DMS dataset sampling and labeling

A broader limitation, both in this study and across the antibody repertoire field, lies in the scarcity and heterogeneity of antigen-binding labeled datasets. Deep mutational scanning (DMS) assays, while powerful, sample only a narrow fraction of sequence space and employ enrichment strategies that bias toward known or likely binders in order to increase the total number of non-negative data^48,50,51,78,79,114^. These enrichment procedures differ across datasets (e.g., mutations of known binders versus high-PLM-likelihood variants), directly affecting the observed local embedding structure independently of the embedding model itself. Such biases could be responsible for making baseline sequence spaces appear to perform comparably or better than PLM embeddings in Vicinity Score and FEF analyses (Fig. 4), as similar sequences are more likely to be sampled.

Additionally, binding classification criteria are inconsistent across studies: some datasets provide discrete labels, while others report continuous affinity values that must be discretized into binding classes (see *Methods*). These thresholds are assay- and antigen-dependent, and although excluding ambiguous intermediate binders reduces misclassification, it can discard informative variants^115^. For instance, influenza HA-H1 specific variants^78^ all bind with varying affinities, whereas in the HER2 Chinery dataset^50^ the “low-affinity binders” are comparable to non-binders. Such inconsistencies and non-standardized affinity measurements impede generalization across studies.

## Methods

### 1. Data acquisition & processing

#### 1.1 iReceptor dataset

A total of 616’095’579 BCR sequences were downloaded from the iReceptor database to serve as background Atlas data. Briefly, the BCR data were filtered based on two conditions: the Organism (ontology ID) was Homo sapiens (NCBITAXON:9606), and the PCR target was IGH. All repertoires, along with study metadata information, were manually downloaded by Study ID. Based on computational feasibility and practicality, we selected 800’000 human HCDR3 sequences for a quantitative assessment of global similarity in PLM embedding distributions (Fig. 2) and 2’151’099 human HCDR3 sequences as a background for plotting antigen-specific datasets using the t-SNE algorithm (Fig. 3). All HCDR3 regions were flanked (i.e., extended at the beginning and end) by the amino acids C (cysteine) and W (tryptophan) to mimic typical antibody structural boundaries.

#### 1.2 Wang dataset

For experiments with W2 distance evaluation, we used 800’000 HCDR3 sequences from an already pre-processed dataset of the study^13^.

#### 1.3 OTS data

5’913’192 pre-processed paired sequences were downloaded from the OTS database^46^ for conducting a quantitative assessment of global similarity in PLM embedding distributions on a TCR data. We sampled at most 399’990 TCRβ sequences for this experiment.

#### 1.4 Randomly generated datasets as a baseline

We also created three random datasets: *random_aa*, *random_shuffled*, *random*. All datasets were generated using NumPy and contain 800’000 sequences.

*Random_aa* is a dataset in which the positional amino acid frequencies in each sequence match those in the experimental iReceptor dataset for HCDR3 sequences. As a result, *random_aa* consists of randomly generated amino acid sequences that replicate both the length distribution and the positional amino acid frequencies observed in the iReceptor dataset.

*Random_shuffled* contains sequences from the iReceptor dataset in which the amino acids within each individual sequence have been randomly shuffled. Consequently, *random_shuffled* consists of sequences with the same length and total amino acid frequency distribution as the iReceptor dataset but disrupted internal structure.

*Random* was constructed from uniformly sampled amino acids, with the same sequence length distribution as in the iReceptor dataset, but different total amino acid frequency.

#### 1.5 Briney dataset

Annotated data was downloaded from Great Repertoire Project GitHub repository (https://github.com/brineylab/grp-paper)^49^ for testing how PLMs can differentiate different donors based on their repertoire information. This dataset contains repertoires taken from 10 healthy donors with varying characteristics. Each participant has 18 samples, consisting of 6 biological replicates (distinct PBMC aliquots), while each biological replicate has 3 technical replicates (independent library obtained from the same RNA) with approximately 3 billion antibody sequences in total. For our experiments, we randomly subsampled at most 18’000’000 HCDR3 sequences from 180 batches.

#### 1.6 Antigen-specific natural antibody sequences dataset

For making an antigen-specific antibody dataset, we downloaded sequences specific to 6 different diseases – COVID-19, influenza, malaria, Ebola, HIV and celiac disease. For SARS-CoV-2 RBD sequences, we selected sequences from the CovabDab database^48^, filtered for “S; RBD” (7’235 unique sequences). Sequences, specific to HIV and influenza proteins, were downloaded from the IEDB database^47^. The dataset contained 171 unique influenza-specific full chains and 284 unique HIV-specific full chains. For Ebola-specific sequences, we downloaded 10’313 unique full-chain sequences from the article^64^. Malaria-specific sequences (1’286 sequences^63^), some influenza-specific sequences (1076 sequences from^63^ and 171 from IEDB^47^) and sequences specific to transglutaminase (TG2) (1’454 unique sequences) – main autoantigen in celiac disease, were provided to us by collaborators^65^. When the information about V, D or J genes was not present in metadata, we aligned sequences using IgBlast ^116^.

#### 1.7 SoNNiA-generated dataset

For comparing antigen-specific datasets using W_2_ distance, we generated a simulated dataset of 1’000’000 heavy chain sequences using the soNNia algorithm, as described in *Issachini et al.*^62^.

#### 1.8 In vitro deep mutational antibody scanning datasets

This study used several deep mutational scanning (DMS) datasets of antibody variants targeting diverse antigens.

Two datasets derived from Trastuzumab variants binding the HER2 antigen were used. The first, described by Chinery et al.^50^, comprises single-chain variable fragment (scFv) variants with mutations restricted to the HCDR3 region (IMGT positions 107–116). These mutations modulate binding affinity toward HER2 and are categorized into three affinity groups: high, medium, and low. To ensure data quality, singleton sequences (clone count = 1), duplicates appearing across multiple affinity groups, and medium binders were excluded, resulting in 172’955 unique high-affinity and 166’527 unique low-affinity variants. The second Trastuzumab dataset (“HER2 Porebski”)^51^ includes variants mutated across 21 amino acids of HCDR3 (24’970 sequences). Preprocessing of the DMS data and assignment of binding-affinity classes followed the pipeline described in Ursu et al^115^. We used the *intensity 8* fluorescence measurement from the original study as a proxy for binding affinity, where higher values indicate stronger binding. Following the original thresholds, sequences were first grouped as binders (intensity 8 > 150; 2’638 sequences), weak binders (122 < intensity 8 < 150; 3’574 sequences), and non-binders (intensity 8 < 122; 18’578 sequences). For downstream analyses, the weak and non-binder categories were merged into a single low-binder class.

Additional datasets were used for Vicinity Score evaluation. These included (i) CR9114 broadly neutralizing antibody variants binding the conserved stem epitope of influenza hemagglutinin (HA) and mutated across the full heavy chain (69’676 sequences)^78^, with high and low-affinity binders separated using a threshold of 9 – log Kd (Fig. S14), and (ii) antibody heavy-chain sequences specific to the HR2 domain of the SARS-CoV-2 spike protein, obtained from the AlphaSeq study^79^ (65’619 sequences).

#### 1.9 Alphaseq study high-low-affinity binders preprocessing

The Alphaseq dataset used for the Vicinity Score analysis (see Fig. 4), consists of predicted affinity measurements of antibody variants given their sequence abundances to the SARS-CoV-2 spike HR2 region^79^ where each variant is defined as a “Protein of Interest” (POI). In two non-overlapping assays, POIs were assayed against the target of interest (“MIT_Target” column, representing the HR2 region of SARS-CoV-2) and three distinct negative controls. Both target and negative control affinities were measured in triplicate. Missing predicted affinity values were replaced with the median value calculated across the negative control replicates according to the original author’s guidelines^79^. Target and negative control replicates were averaged separately to yield one average target affinity and one negative control affinity for each POI which was used to classify them into High and Low-affinity binders. All the above data preprocessing steps were conducted separately for each assay to accommodate differences in technical variability.

POIs were classified as a High-affinity binder if they satisfied three conditions: (1) The POI has at most one missing predicted affinity value, (2) their average target affinity was lower (i.e. stronger binding) than the binding threshold (5th percentile of the average negative control affinity), and (3) the difference between the average negative control affinity and the average target affinity was less than -1 nM log10 KD, indicating at least 10 times stronger binding to target than to the negative control. POIs were classified as “Low-affinity binders” if (1) there were less than or equal to 2 missing replicates (out of three) in predicted target affinity, and (2) their binding affinity (KD) was lower than the binding threshold (see above) **or** the difference between the average negative control affinity and the target affinity is below -1 nM log10 KD (Fig. S15). All other POIs were discarded. In total, out of 104’972 entries, 16’501 POIs were classified as High-affinity binders and 49’118 as low-affinity binders.

## 2. Generation of antibody sequence embeddings

### 2.1 Language models and baseline used in the study and different preprocessing

Amino acid sequence embeddings and attention matrices were extracted from the final and intermediate embedding layers of ESM2 and an antibody-specific language model, AntiBERTa2-CSSP. Token-level embeddings were then pooled into sequence-level embeddings by averaging all token-level embeddings of a sequence. For a ‘baseline’ comparison, we calculated one-hot encoded (OHE) representations using the ‘OneHotEncoder’ from the immuneML python library^117^, where each amino acid is represented by a one-dimensional vector, corresponding to the respective standard amino acid.

#### 2.1.1 AntiBERTa2

AntiBERTa2-CSSP^45^ (AB2) is an antibody-specific transformer model based on the RoFormer architecture, a bi-directional transformer encoder model with rotary position embeddings^44,118^. AB2 is pre-trained in two stages. The first stage consists of unsupervised pre-training on ∼779.4 million human antibody sequences with a masked language modeling (MLM) objective. In the second step, AB2 is trained on 1237 antibody structures via *contrastive sequence-structure pre-training* (CSSP) using ESM-IF as a pre-trained structure encoder with frozen weights. The resulting model has ∼202 million parameters, 16 embedding layers with 1024 embedding dimensions.

#### 2.1.2 ESM2

“esm2_t33_650M_UR50D” (ESM2) is a member of the ESM-2 PLM family^44^ and has been trained on over ∼65 million unique protein sequences present in the UniRef database^119^. Its neural network has a total of ∼650 million parameters and contains 33 transformer layers with 1280 embedding dimensions.

### 2.2 Visualization methods used in the study

#### 2.2.1 t-SNE

T-distributed stochastic neighbor embedding (t-SNE) algorithm is a nonlinear dimension reduction and data visualization method^120^. We used the openTSNE^121^ package (v1.0.2) and main parameters used for fitting were: perplexity=30, metric="cosine", random_state=123, early_exaggeration=25.0, initialization="pca".

## 3. Global sequence analysis

### 3.1 Approximation of Wasserstein-2 distance

The Wasserstein-2 distance (W₂) was calculated as the distance d(.) between the normal distribution p_w_(.) with mean m_w_ and covariance C_w_ and between the normal distribution p(.) with mean m and covariance C^55^:

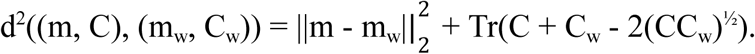

In our setting, the input to W₂ is the set of repertoire embeddings for each group. From these embeddings, we compute the empirical mean vector and covariance matrix that define a Gaussian approximation of the distribution. These parameters are then used to compute the Wasserstein-2 distance between the two Gaussians, yielding a single scalar distance value. Although PLM embeddings are not perfectly Gaussian, this approximation is widely used because it is computationally efficient and captures meaningful first- and second-order differences at scale^55,84,85^. For the Python implementation, scikit-learn (v1.3.0) package was used.

### 3.2 Distinguishing of donor-specific repertoires using W_2_ distance

In this experiment, we utilized different modifications of the *Briney et al.*^49^ dataset to explore the applicability boundaries of the respective method. Specifically, we used the following modifications. The *standard* Briney et al. dataset – from which a maximum of 100’000 random sequences of HCDR3 regions were selected. In the *flanked dataset*, for each selected HCDR3 region, a cysteine (C) was added to the beginning if this amino acid was not present at the start of the region, and a tryptophan (W) was added to the end if it was not present at the end of the sequence. This way, we obtained HCDR3 regions with uniform beginnings and endings. For the *length-filtered dataset*, only HCDR3 regions of 15 amino acids were selected. In the *length filtering + flanking dataset*, the two modifications described above were applied simultaneously. For the *unique sequences + length filtering* dataset, from the length-filtered HCDR3 regions, only unique sequences were retained (only one occurrence of each region was allowed in the dataset). In *clonal filtering within a single donor* dataset, for each donor, the clonality of all HCDR3 regions was determined by HILARy^122^ (v1.0.13), followed by cloning filtering within the same donor. Finally, for *single linkage clustering across all donors* dataset, single linkage clustering was performed for all HCDR3 regions across all donors by applying HILARy with fixed threshold for Hamming distances in each group, followed by the filtering of each cluster of similar sequences so that only one sequence remains out of every group.

For the first four dataset modifications (the standard dataset, flanked, filtered by length and filtered by length + flanked), we used all 180 available replicates from the *Briney et al.* dataset. To ensure accurate computation of the W2 distance, we set a minimum batch size of 40’000 sequences, while for computational feasibility, we imposed an upper limit of 100’000 sequences.

For subsequent modifications, we selected only replicates with enough sequences to compute exact W_2_ metric. For unique sequences + length filtering, we used 100 replicates, for clonal filtering within a single donor modification – 117 replicates and for single linkage clustering across all donors – 83 replicates.

For filtered replicates, we calculated the embeddings according to different encodings (in our experiment, ESM2^44^, AntiBerta2^45^, OHE and left-right model^61^)and calculated the pairwise W_2_ distance for each replicate pair using code from the study^55^. Next, we applied the agglomerative clustering algorithm (scikit-learn v1.3.0) to the pairwise W_2_ distance matrix, with a fixed number of expected clusters equal to the number of different donors (by default, 10). To evaluate the clustering quality, we used an Adjusted Rand Index (where 1 means perfect overlap). We used the adjusted_rand_score function from the scikit-learn (v1.3.0) package^75^. For Jaccard index computation, no sequence encoding was required; instead, Jaccard index matrices were obtained directly using a custom Python implementation.

## 4. Visual comparison of simulated, natural and antigen-specific sequence embeddings

### 4.1 Comparison of antigen-specificity labeled datasets

Each group of antigen-specific sequences (HCDR3 only) was embedded using AB2 and ESM2, and then mean pooling was applied. We then computed the Wasserstein-2 (W2) distance between the embeddings of each pair of antigen-specific groups. Within-group variability was assessed by splitting each group randomly in half and computing the W2 distance between the two halves. This procedure was repeated 10 times with different random splits.

For the iReceptor and simulated groups, which were orders of magnitude larger than the others, we downsampled to 20,000 sequences each to maintain comparability.

### 4.2 Dimensionality reduction with T-SNE

T-SNE was applied to reduce the dimensionality of the embedding space. A t-SNE model was first fitted on the 2’151’099 iReceptor embeddings (mean-pooled, HCDR3-only sequences flanked by C–W) using the .fit() function (parameters used: perplexity=30, metric="cosine", random_state=123, early_exaggeration=25.0, initialization="pca"). After model fitting, both the iReceptor and antigen-specific embeddings were projected into the learned low-dimensional space using the .transform() function, following the guidelines provided in the openTSNE^121^ documentation. This procedure yielded two-dimensional coordinates (t-SNE 1 and t-SNE 2) for downstream analyses and visualization.

### 4.3 Relative density distributions

To quantify density distributions for each combination of dataset and binding label in the t-SNE-reduced space, we implemented a kernel density estimation (KDE)-based approach. First, we determined the overall range of t-SNE coordinates along the first and second dimensions, across all antigen groups and iReceptor, to ensure shared coordinate range and comparability across groups. Then densities were estimated for each group separately, using a two-dimensional KDE (implemented via the kde2d function from the MASS^123^ package in R), evaluated on a uniform grid of 100 × 100 points across the defined ranges. The resulting density estimates were scaled by dividing each value by the maximum density within the group, yielding a density score between 0 and 1. Group-wise densities were then combined for downstream visualization and analysis.

### 4.4 Density difference calculation

We followed the same procedure as the relative density difference distribution, with just two small differences. To compute the shared coordinates range, the background was also included. To obtain the density difference value, the density for each group was normalized so that its total integrated density (area under the surface) equaled 1. Subsequently, the resulting normalized densities were subtracted between groups (group 2 - group 1).

## 5. Local functional clustering

The Vicinity Score measures the degree to which AIR sequences with the same functional label are spatially homogeneous in PLM embedding space at a given radius R. For each AIR, precision is defined as the fraction of sequences within a specified cosine distance radius that share its label. The Vicinity Score is the median of these precision values across all AIRs at a given R, providing a radius-dependent measure of functional local spatial homogeneity. To account for biases due to uneven densities, we included a resampling step in which Precision values are sampled inversely by their local density before the Vicinity Score calculation.

The first step to compute the Vicinity Score is to calculate the Precisions for each AIR, at a given cosine distance R. To do so, we utilize the K-nearest neighbors (KNN) algorithm from sckit-learn^75^ to determine the distances from each AIR to its neighbors (restricted to the first 2000th to address computational constraints).

To define the set of R values, we selected 15 evenly spaced thresholds spanning the range of pairwise distances among KNNs. For each R, we computed the precision of every AIR, defined as the fraction of neighboring sequences within that radius that share its binding label. The Vicinity Score at a given R is then obtained as the median of these Precision values across all AIRs. As absolute cosine distances, which determine the radii (R), are not directly comparable across embedding spaces, we mapped these radii to a unified scale termed Neighborhood size, defined as the average number of nearest neighbors (K) at each radius R.

Sparsely populated regions in the embedding space can lead to high variance in Vicinity Scores due to the limited number of neighboring sequences. To counteract this issue, we applied a weighted inverse sampling approach to the precision values used to compute the Vicinity Scores at each R. The inverse sampling weighted by local density role (i.e., precision values are resampled with probability inversely proportional to local density) is to reduce the overrepresentation of Vicinity Scores coming from densely populated regions. The local density of each sequence in embedding space is approximated by its corresponding embedding proximity graph (EPG) node degree (see section *Topological comparisons with proximity graphs*). For each R, we performed inverse sampling based on local density, selecting 25% of the total precision values.

When Vicinity Scores are computed in sequence space rather than embedding space, the procedure remains identical, with two modifications. First, distances are calculated using Levenshtein distance (LD) instead of cosine similarity, and radii R are defined accordingly. Second, for the inverse weighted sampling step, the local density is approximated by the sequence proximity graph (SPG) node degree (see Topological comparisons with proximity graphs). Of note, we included only non-redundant sequences within each dataset in all Vicinity analyses to avoid the effect of identical sequences on the outcome.

### 5.1 Dataset input and embeddings complexity approaches

We evaluated five input configurations with varying degrees of contextual antibody sequence information.

- All HCDR: Representations of amino acid sequences from heavy chain CDRs. HCDR1, HCDR2 and HCDR3 were concatenated into a single vector via separator tokens. (AB2 used [SEP] token, ESM2 used the token) inspired by *Briney et al*^34^.
- HCDR3 only: Representations of amino acid sequences from heavy chain CDR3.
- Heavy chain: Representations of amino acid sequences from heavy chain variable domains (VH).
- HCDR3 Extracted: Representations of V_H_ amino acid sequences that were truncated to retain only values at HCDR3 positions.
- Paired chain: Representations of paired antibodies. Amino acid sequences from heavy-and light chain variable domains (V_H_ and V_L_) were concatenated into a single vector via separator tokens (see All HCDR).

Of note, we performed HCDR3 sequence flanking by adding a single ‘C’ residue to the start and a single ‘W’ residue to the end of the sequence, unless they were already present.

To assess the effect of embedding dimensionality and representation on the vicinity score, three embedding modes were extracted from each hidden layer of the models: (i) Mean pooling represents the current state-of-the-art approach to reduce the dimensionality of the output of a given layer by averaging the output values along the amino acid (residue) dimension, leading to a 1 dimensional vector per residue as embedding. To preserve more amino acid token information, we mask all embeddings of special tokens (pad, beginning-/end-of-sequence, separators, etc.) and perform the mean calculation only on amino acid token embeddings. (ii) Attention matrices represent extracting the attention matrices for all the attention heads from a given layer, and averaging them. This returns a L x L matrix (where L = number of unmasked tokens in context window), which is then flattened to a 1-dimensional vector of length **L²**. (iii) Unpooled embeddings represent just getting the raw high-dimensional output from the model, and not applying any dimensionality reduction techniques, but just flattening the whole matrix into a 1-dimensional vector of size L x E (**E** represents the Embedding dimensionality of the model).

All representations were extracted with PEPE^124^, as it requires only a single forward pass per PLM and input configuration to extract multiple embedding modes from all hidden layers.

### 5.2 Topological comparisons with proximity graphs

To quantitatively approximate the topology of a representation space, we construct unweighted, undirected proximity graphs in which each node represents a sequence and edges connect adjacent nodes. In a sequence proximity graph (SPG) two nodes are considered adjacent if their sequences are 1 LD apart. An embedding proximity graph (EPG) is a graph where two nodes are considered adjacent if the cosine distance between their embeddings is below a threshold **D_thr_**. We define D_thr_ for each embedding space as the average cosine distance between embeddings whose sequences are 1 LD apart. For computational feasibility, the distance calculation is limited to the 1000 nearest neighbors of each node.

## Code availability

The code used to extract PLM embeddings and attention matrices is available at https://github.com/csi-greifflab/pepe-cli. Scripts to reproduce our experiments are available from https://github.com/csi-greifflab/airr_atlas.

## Data availability

Every dataset used can be accessed by its related publication, mentioned in Methods (Data acquisition & processing section). The dataset information can be accessed in the Supp. Table 1.

## Acknowledgements

We thank Philippe A. Robert for helpful comments that led to increased readability of the manuscript.

## Supplementary Tables

**Supp. Table 1:** | Information about public datasets used in this study. The name of the dataset is the same as in Methods.

**Supp. Table 2**: | Information about the effect sizes between the Vicinity score curves showed in Fig. 4.

---

**Supplementary figure 1:**
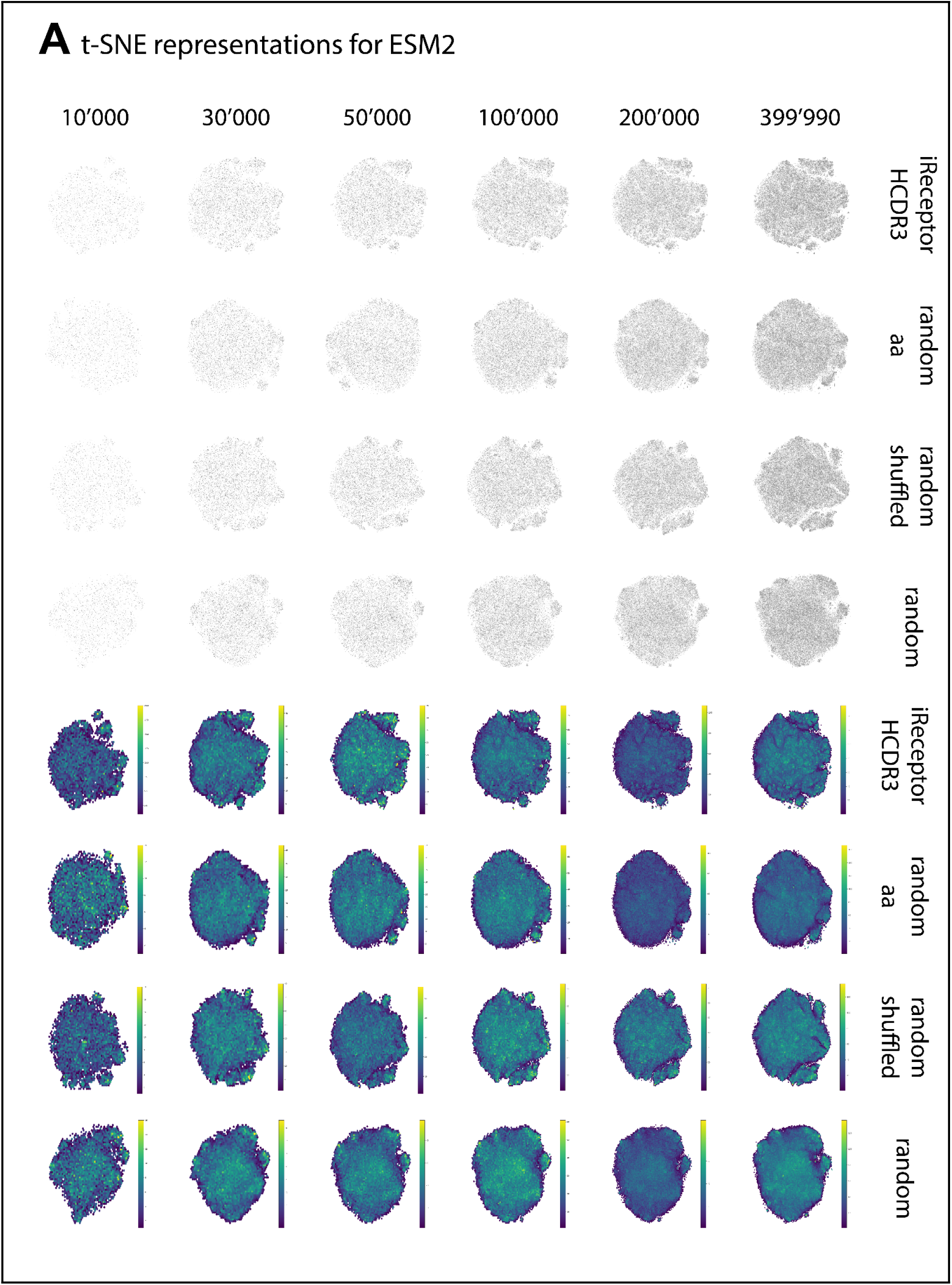
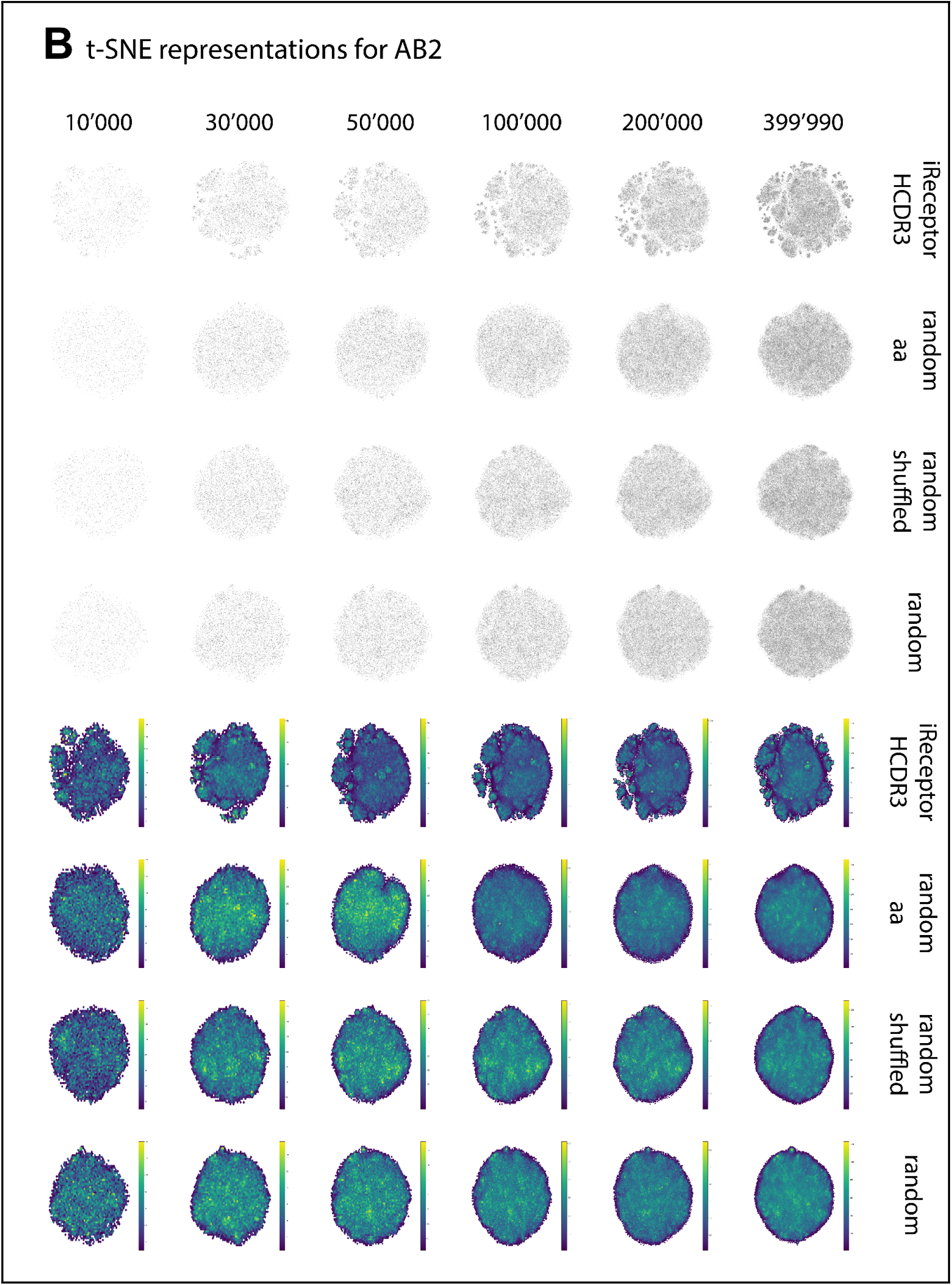
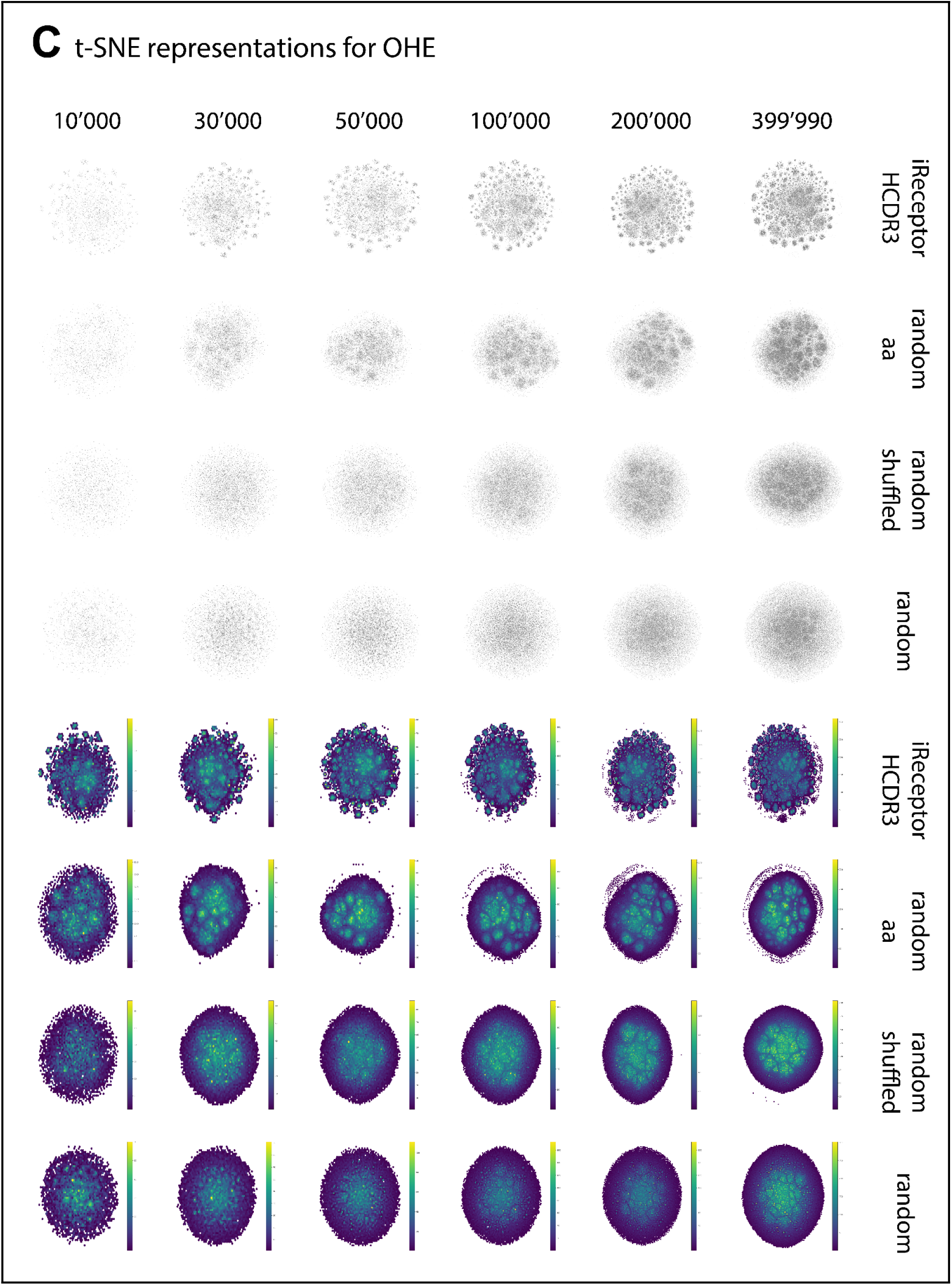
PLM embeddings of HCDR3 BCR sequences reveal distinct high-dimensional structure starting from 30,000 sequences. Each plot represents a t-SNE visualization (left) and a hexbin density plot (right) of the A) AB2, B) ESM2, and C) OHE embeddings, with batch sizes noted above the plots and the corresponding model labeled on the right. Experimental iReceptor data form well-defined topologies, while randomly generated controls (*random_aa*, *random_shuffled*, *random*) lack visible structure, suggesting that AB2, ESM2, and OHE embeddings capture biologically meaningful patterns absent in randomized sequences. **Refers to main figure:** Fig. 2

**Supplementary figure 2:**
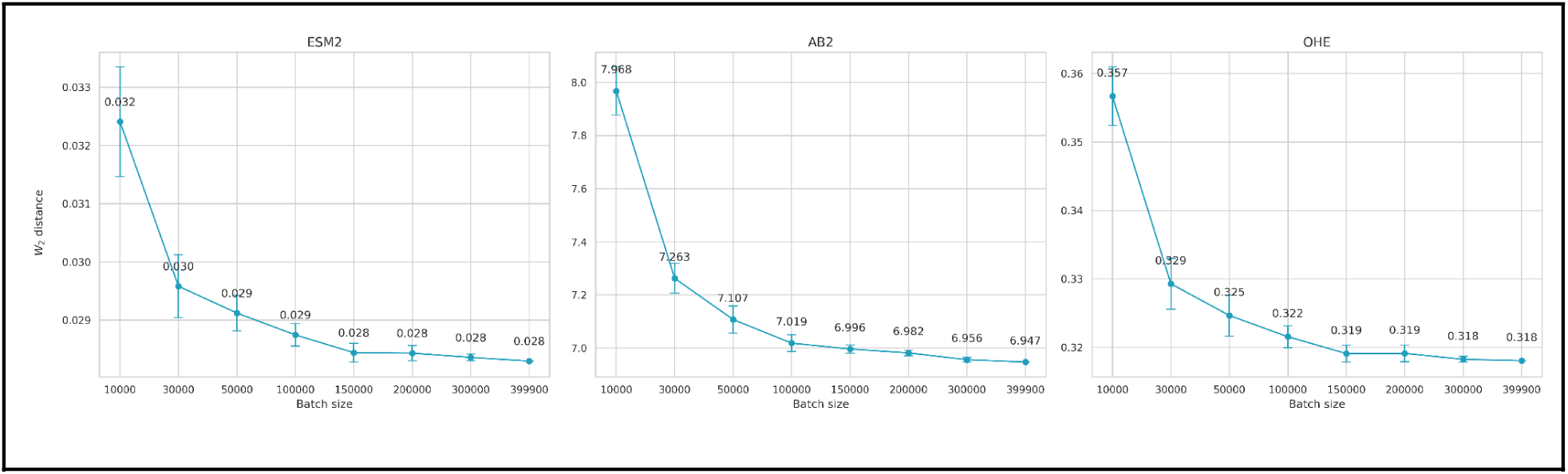
Accurate estimation of mean and covariance for TCRβ embeddings using AB2, ESM2, and OHE requires larger sample sizes compared to BCRs. Each plot displays a boxplot of pairwise W₂ distances for each batch size (x-axis) compared to randomly sampled TCR sequences totaling 399’990 from the iReceptor database. For every batch size tested, the W₂ distances for TCRs remain consistently higher than those observed for BCRs, indicating less compact embedding distributions. Each pairwise distance was calculated ten times to demonstrate that, with a sufficiently large sample, the variance in W₂ estimation becomes minimal. Refers to main figure: Fig. 2

**Supplementary figure 3:**
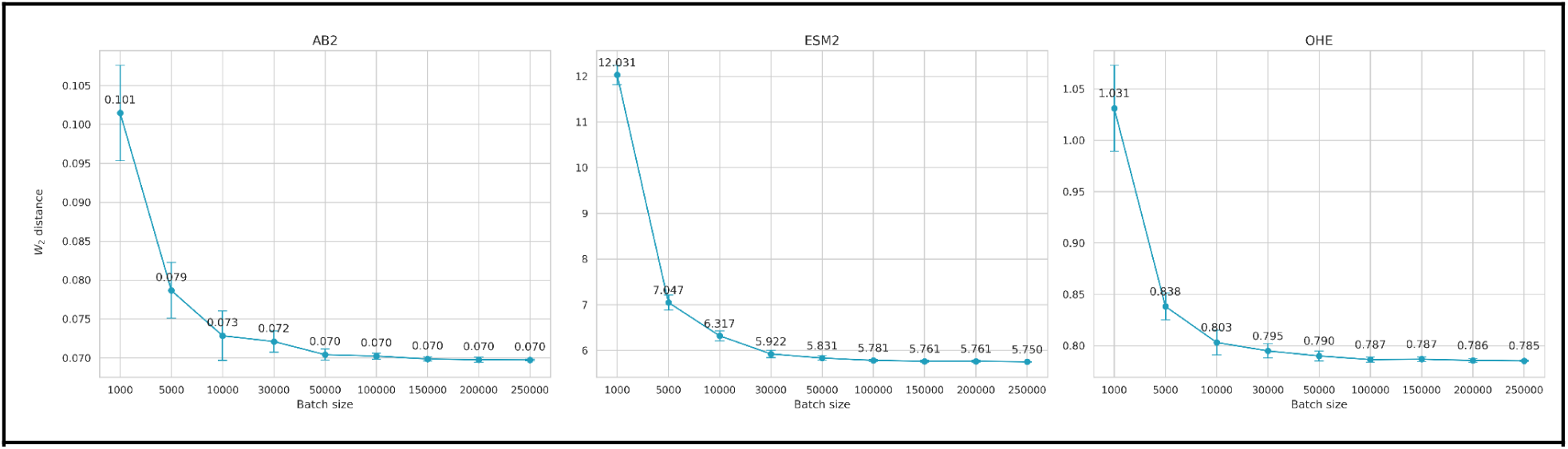
Accurate estimation of mean and covariance for Trastuzumab variant HCDR3 targeting HER2 embeddings using AB2, ESM2, and OHE requires larger sample sizes compared to iReceptor BCRs. Each plot displays a boxplot of pairwise W₂ distances for each batch size (x-axis) compared to randomly sampled Trastuzumab variant HCDR3 sequences totaling 250’000 from the Trastuzumab dataset (see Methods). For every batch size tested, the W₂ distances for Trastuzumab variants remain consistently higher than those observed for iReceptor BCRs, indicating less compact embedding distributions. Each pairwise distance was calculated ten times to demonstrate that, with a sufficiently large sample, the variance in W₂ estimation becomes minimal. Refers to main figure: Fig. 2

**Supplementary figure 4:**
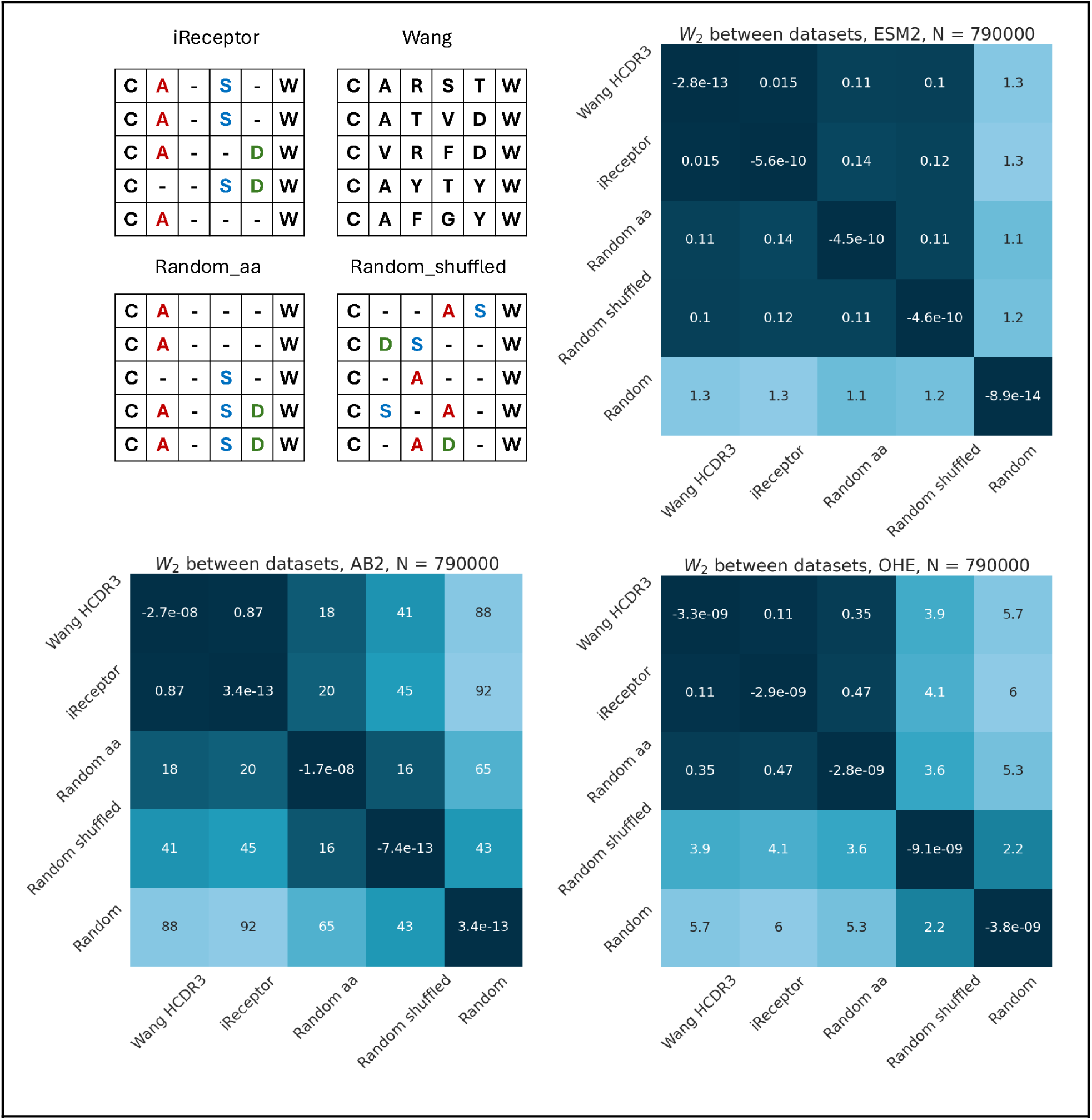
W₂ distance applied to PLM embeddings reveals a clear separation between experimental immune repertoires (Wang and iReceptor) and randomly generated controls, including those matched for sequence length and positional amino acid frequencies. The plots consist of heatmaps depicting pairwise distances among the datasets: Wang, iReceptor, *random_aa* (maintains the frequency distribution at each position similar to experimental data), *random_shuffled* (amino acids in each sequence are shuffled) and *random* (uniform frequency distribution at each position) (see Methods). Heatmaps show that pairwise distances between experimental datasets are consistently smaller than the distances between experimental and random datasets, confirming the ability of W₂ to capture biologically meaningful differences in sequence distributions. Refers to main figure: Fig. 2

**Supplementary figure 5:**
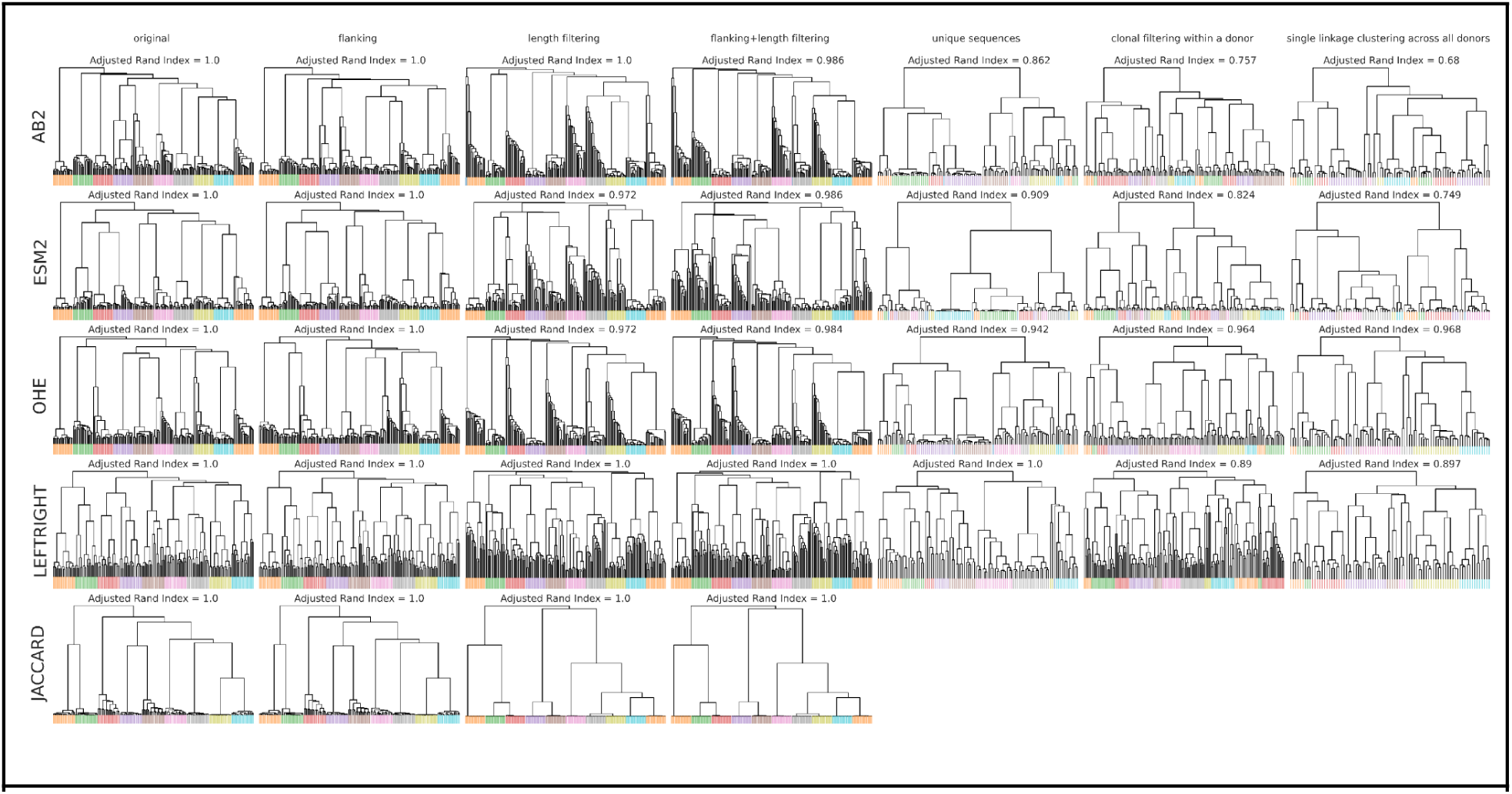
Evaluating the performance of different pipeline modifications in revealing donor-specific repertoire structure. Y-scale is symmetric logarithmic. The different rows stand for different sequence representations. ‘AB2’ – AntiBerta2 embeddings and W_2_ distance calculation, ‘ESM2’ – ESM2 embeddings and W_2_ distance calculation, ‘OHE’ – OHE embeddings and W_2_ distance calculation, ‘LEFTRIGHT’ – SONIA left-right model and Pearson correlation, ‘JACCARD’ – Jaccard index calculation. Different columns stand for different data modifications. ‘ORIGINAL’ – for each replicate, 100 thousand HCDR3 sequences were randomly sampled. ‘FLANKING’ – for each sampled HCDR3 sequence, a single cysteine residue to the start of the sequence and a single tryptophan residue to the end of the sequence was added, unless they were already present. ‘LENGTH FILTERING’ – for each replicate, we sample 100 thousand HCDR3 sequences that are 15 amino acids long. ‘FLANKING + LENGTH FILTERING’ – selected 15-amino-acid-long HCDR3 sequences were flanked with cysteine and tryptophan residues from the start and the end, respectively. ‘UNIQUE SEQUENCES + LENGTH FILTERING’ – from the length-filtered HCDR3 regions, only unique sequences were retained (only one occurrence of HCDR3 was allowed in the dataset). ‘CLONAL FILTERING WITHIN A SINGLE DONOR’– for each donor, the clonality of all HCDR3 regions was determined, followed by clonal filtering within the same donor. ‘SINGLE LINKAGE CLUSTERING ACROSS ALL DONORS’– single linkage clustering was performed for all HCDR3 regions across all donors. Since there are no identical sequences between replicates after unique sequences and clonal family filtering, Jaccard index always equals 0. Therefore, the last three plots in the ‘JACCARD’ row are not shown. **Refers to main figure:** Fig. 2

**Supplementary figure 6:**
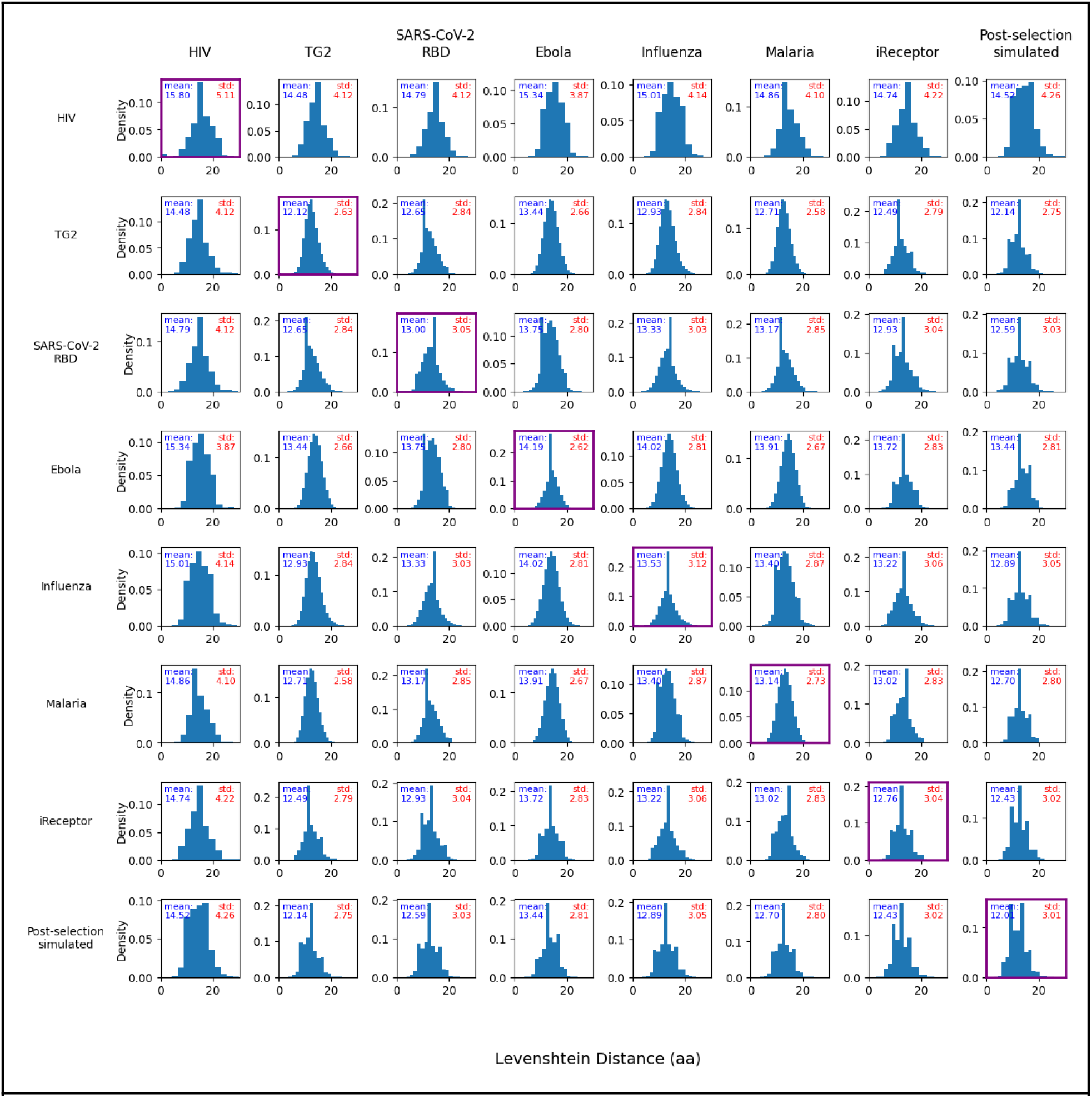
Pairwise Levenshtein distance across HCDR3s within antigen-specific datasets, iReceptor meta-repertoire and simulated datasets. Pairwise Levenshtein (edit) distances were computed between all HCDR3 amino acid sequences within each antigen-specific dataset (diagonal, in purple) and between datasets (off-diagonal). Each panel shows the probability density of pairwise Levenshtein distances and reports the mean (blue) and standard deviation (red). The number of sequences for each antigen matches those shown in Fig. 3. Given computational restrictions, iReceptor and Post-selection simulations were downsampled to 20k sequences each.

**Supplementary figure 7:**
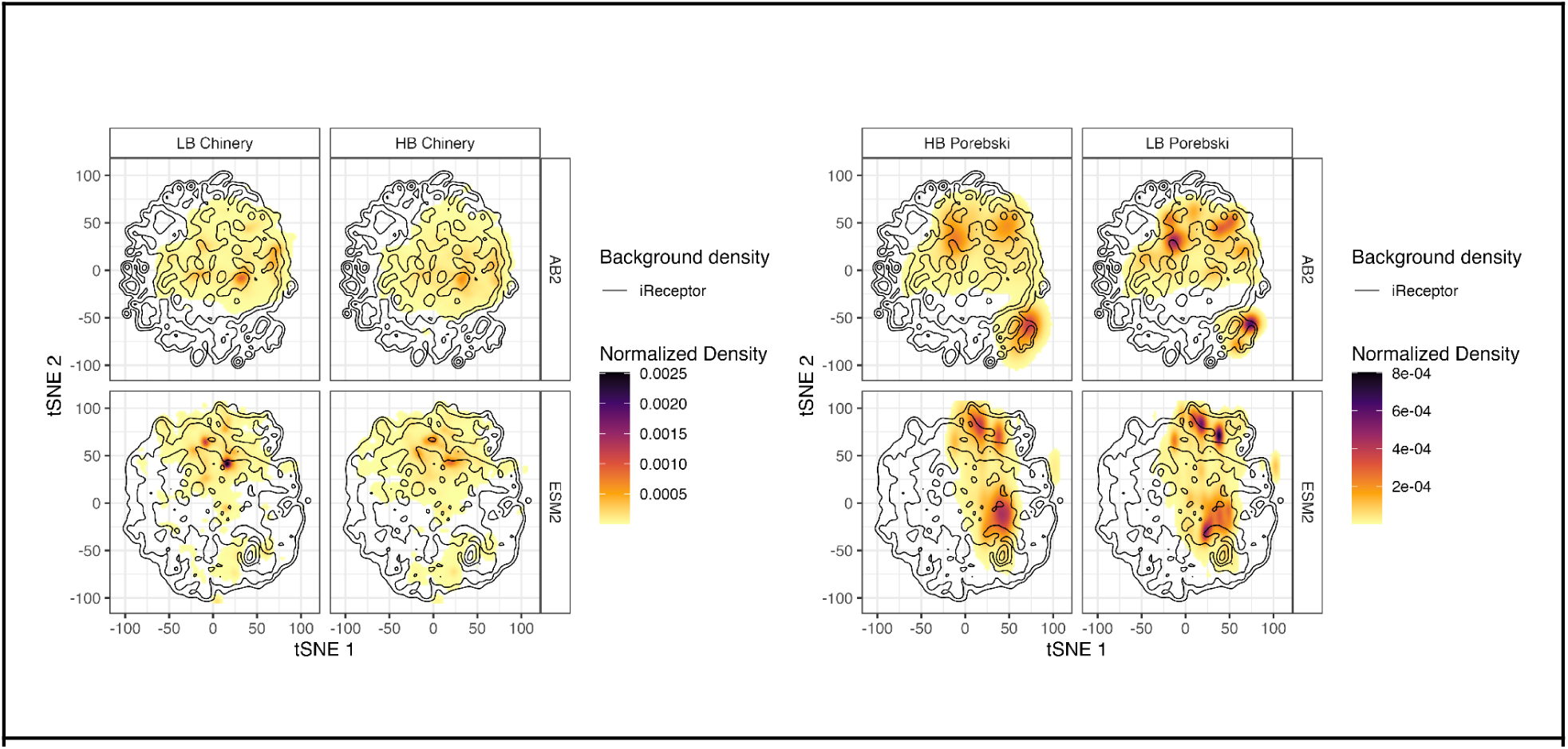
Density distribution of HER2-specific antibodies on top of the two-dimensional projections (t-SNE 1 and t-SNE 2) of the iReceptor background repertoire. The contours represent the density of the background meta-repertoire (iReceptor), overlaid with the density of the HER2-specific antibodies. The antibodies from the two different studies (Chinery et al., Porebski et al.,)^50,51^ were plotted separately as they have different lengths and characteristics (see Methods). Within each study, densities of high (HB) and low-affinity (LB) binders are plotted, with integral normalization (see Methods) performed within each HB/LB set. The HER2-Chinery dataset contains 49,721 HBs and 47,328 LBs, the HER2-Porebski contains 2,638 HBs and 22,152 LBs, and the iReceptor meta-repertoire comprises approximately 2 million sequences. Refers to main figure: Fig. 3

**Supplementary figure 8:**
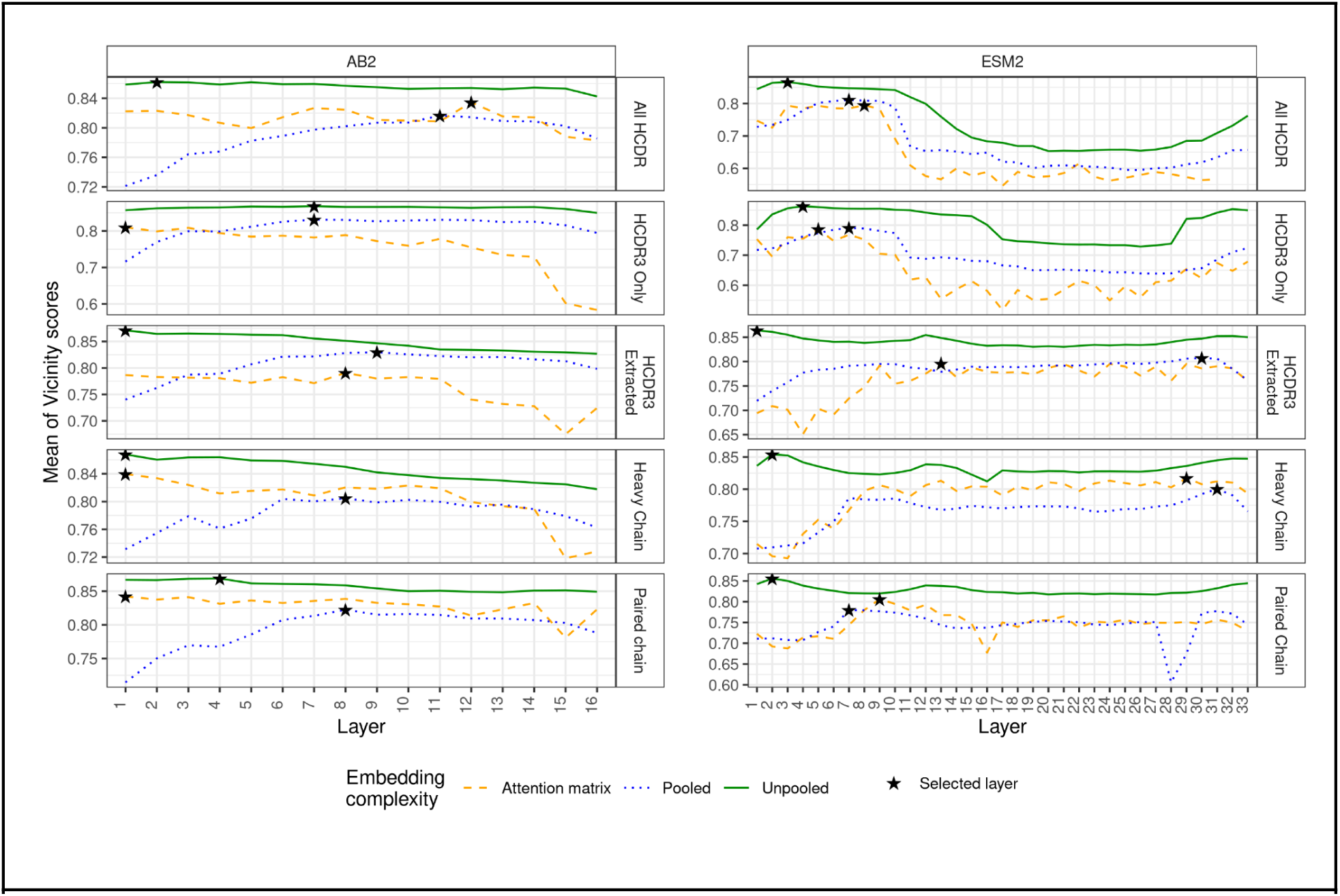
Layer-wise average Vicinity Scores of high-affinity binders in the HER2 Chinery dataset. The plot shows how the mean Vicinity Scores (*m*_c_ ) vary across layer depth. The HER2 Chinery dataset was embedded using different combinations of embedding complexity (represented by line types), sequence content, and models (shown as faceting variables; see Methods). The star indicates the layer with the highest mean Vicinity Score *m*_c_, which was selected for downstream analysis. **Refers to main Figure:** Fig. 4

**Supplementary figure 9:**
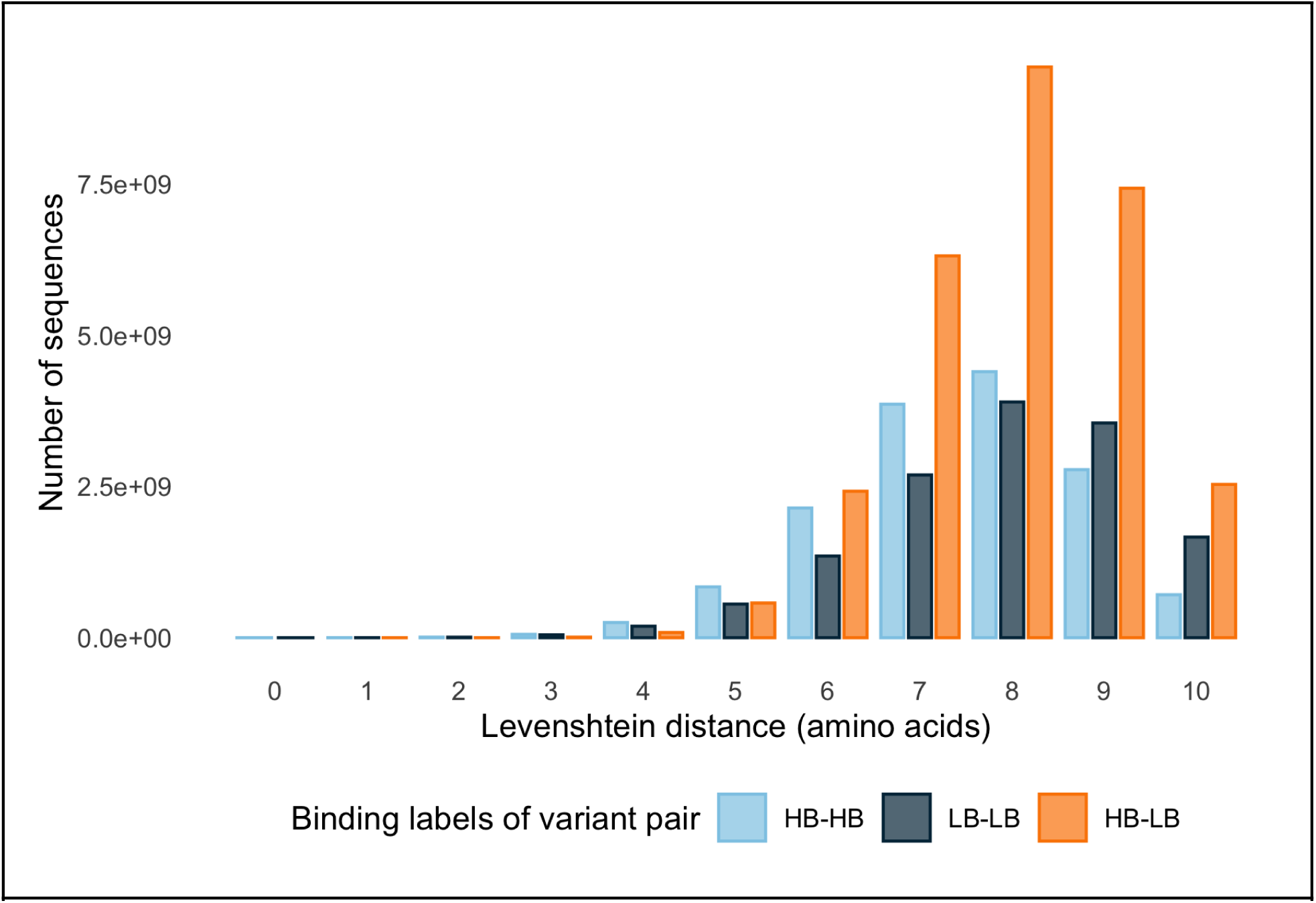
High-affinity and low-affinity binding variants of HER2 are not trivially separated by edit distance. Pairwise edit distance was calculated for 339482 Trastuzumab variants from the HER2-Chinery dataset. Variant pairs were grouped by binding labels (HB-HB, LB-LB and HB-LB) and are indicated by color. The most frequent Levenshtein distance for all binding label groups was 8 amino acids. Abbreviations: HB (high-affinity binder), LB (low-affinity binder). **Refers to main Figure:** Fig. 4

**Supplementary figure 10:**
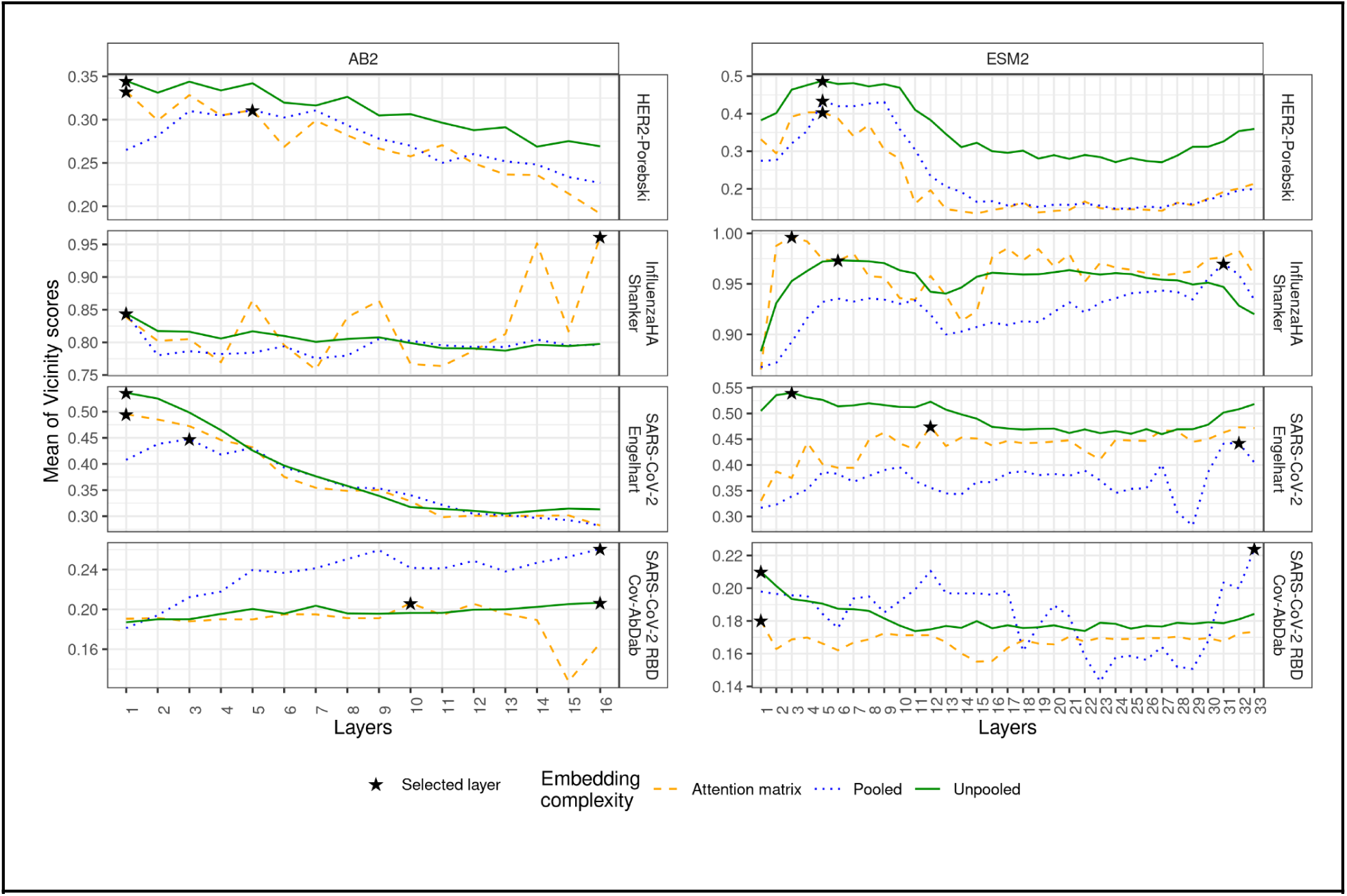
Layer-wise average Vicinity Scores of high-affinity binders across DMS dataset with decreasing positional restriction for mutations. The plot shows how the mean Vicinity Scores (*m_c_* ) vary across layer depth. The selected datasets were embedded using different combinations of embedding complexity (represented by line types), sequence content, and models (shown as faceting variables; see Methods). The star indicates the layer with the highest mean Vicinity Score, which was selected for downstream analysis. Refers to main Figure: Fig. 4

**Supplementary figure 11:**
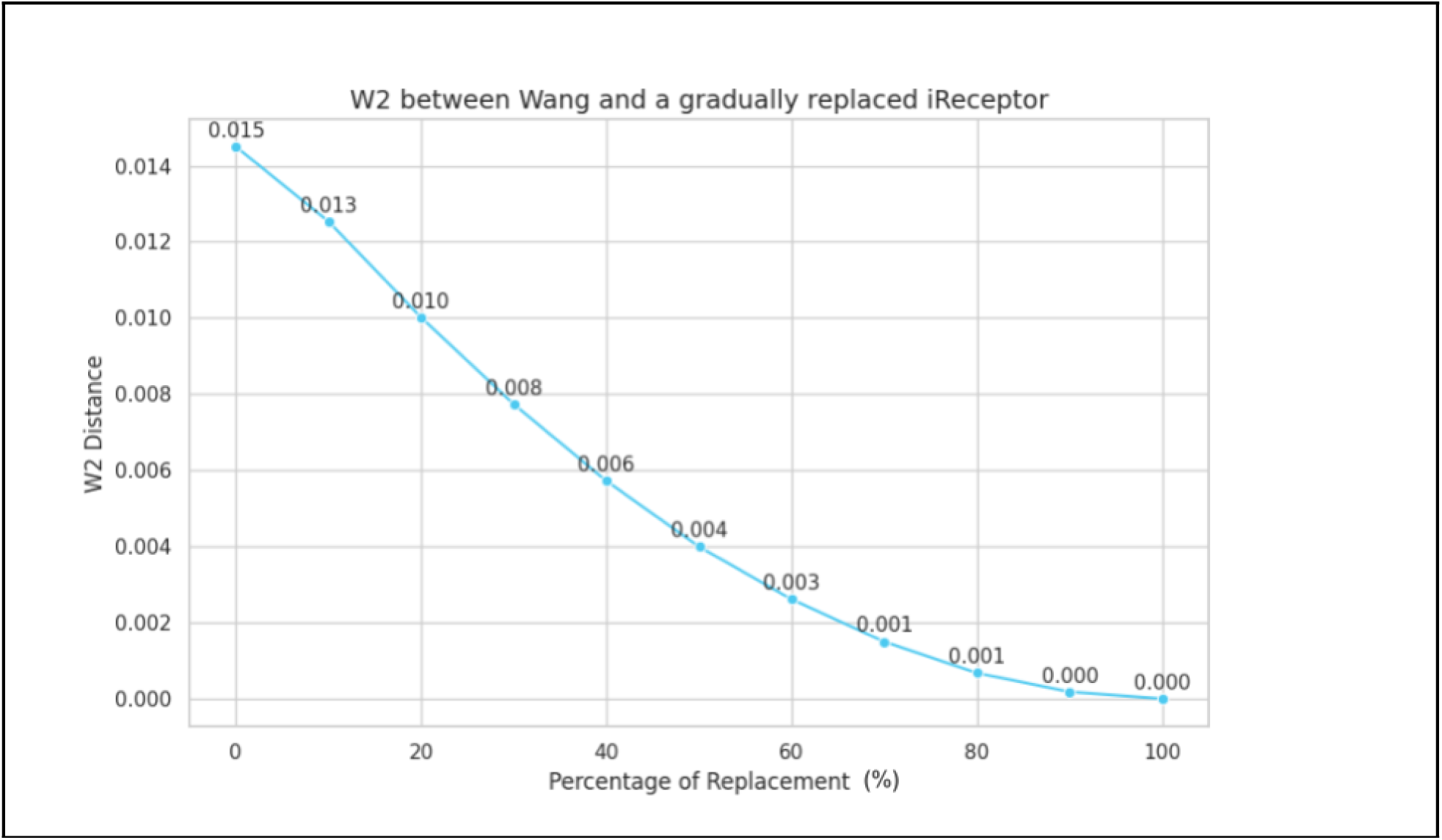
W₂ decreases non-linearly as more sequences are shared between two repertoires. To generate this curve, we constructed controlled mixtures of two datasets (Wang and iReceptor) by gradually replacing a fraction of sequences from one dataset with sequences from the other, and recalculating the pairwise W₂ distance at each replacement step. This illustrates the sensitivity of W₂ to redundancy and public clonotypes: repertoires with extensive overlap appear more similar even when their unique components differ substantially. Consequently, interpretation of W₂ requires accounting for clonality and shared sequence content to avoid over- or underestimating repertoire similarity. **Refers to main figure:** Fig. 2

**Supplementary figure 12:**
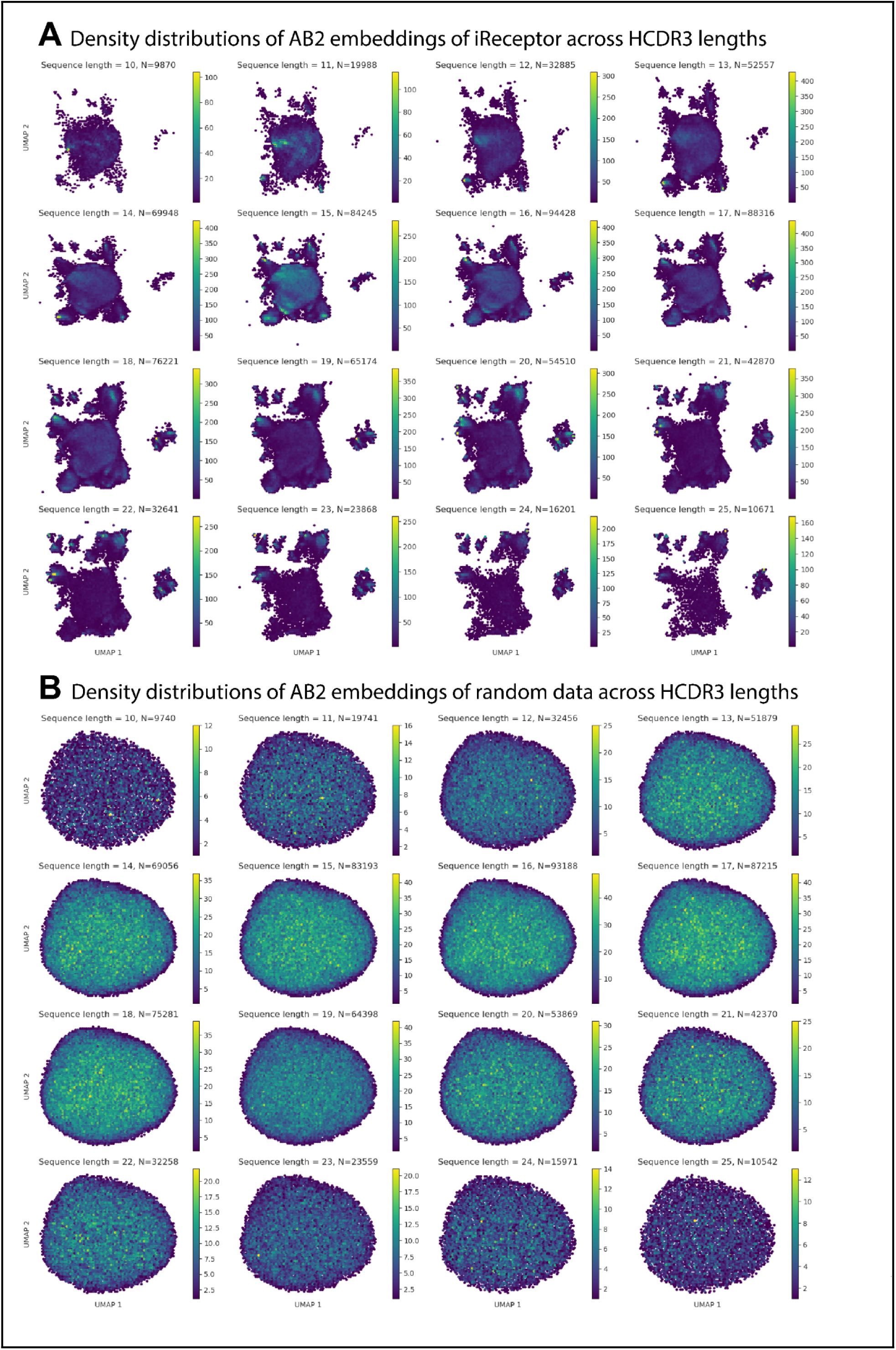
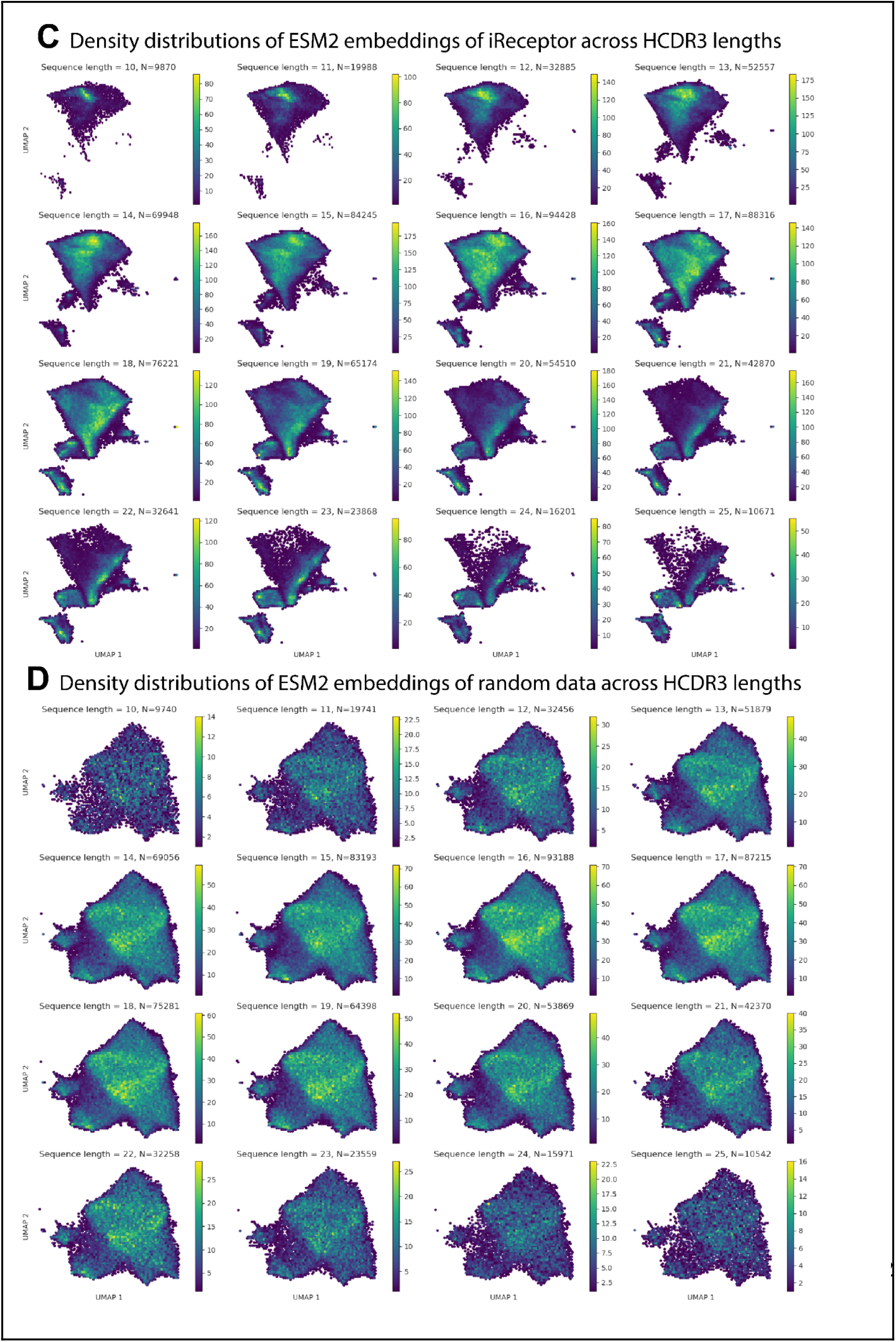
Length bias in PLM embeddings leads to distinct density patterns for sequences of different HCDR3 lengths. A – hexbin density maps of AB2 embeddings for the iReceptor dataset. B – hexbin density maps of AB2 embeddings for the random data. C – hexbin density maps of ESM2 embeddings for the iReceptor dataset. D – hexbin density maps of ESM2 embeddings for the random data. Each subplot corresponds to sequences of a given HCDR3 length. Hexbin coloring reflects local point density. Across both experimental and random datasets, sequences of the same length cluster into distinct regions of embedding space, while distributions for different lengths remain separated. This demonstrates that PLM embeddings are strongly structured by sequence length, independent of repertoire origin. **Refers to main figure:** Fig. 2

**Supplementary figure 13:**
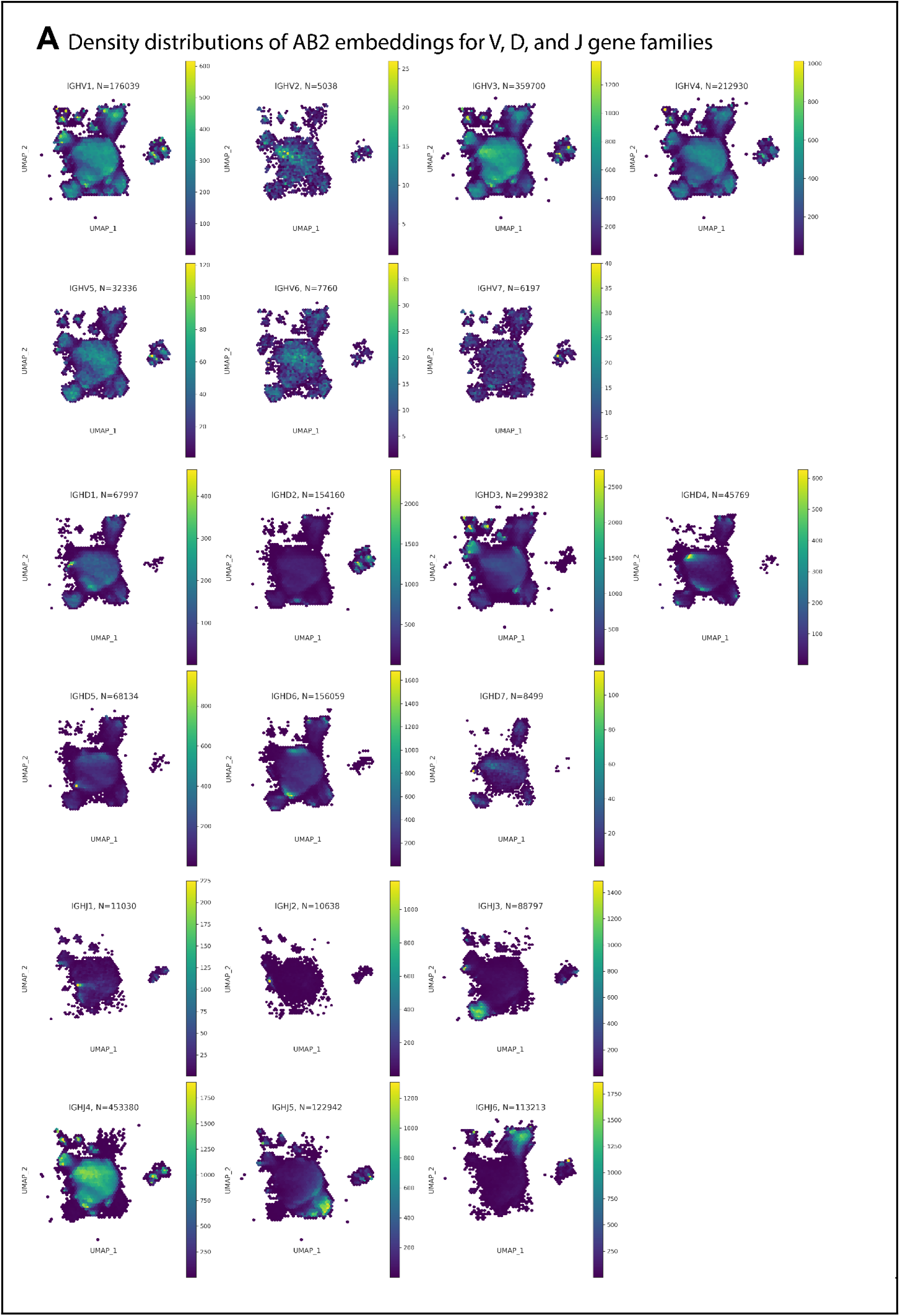
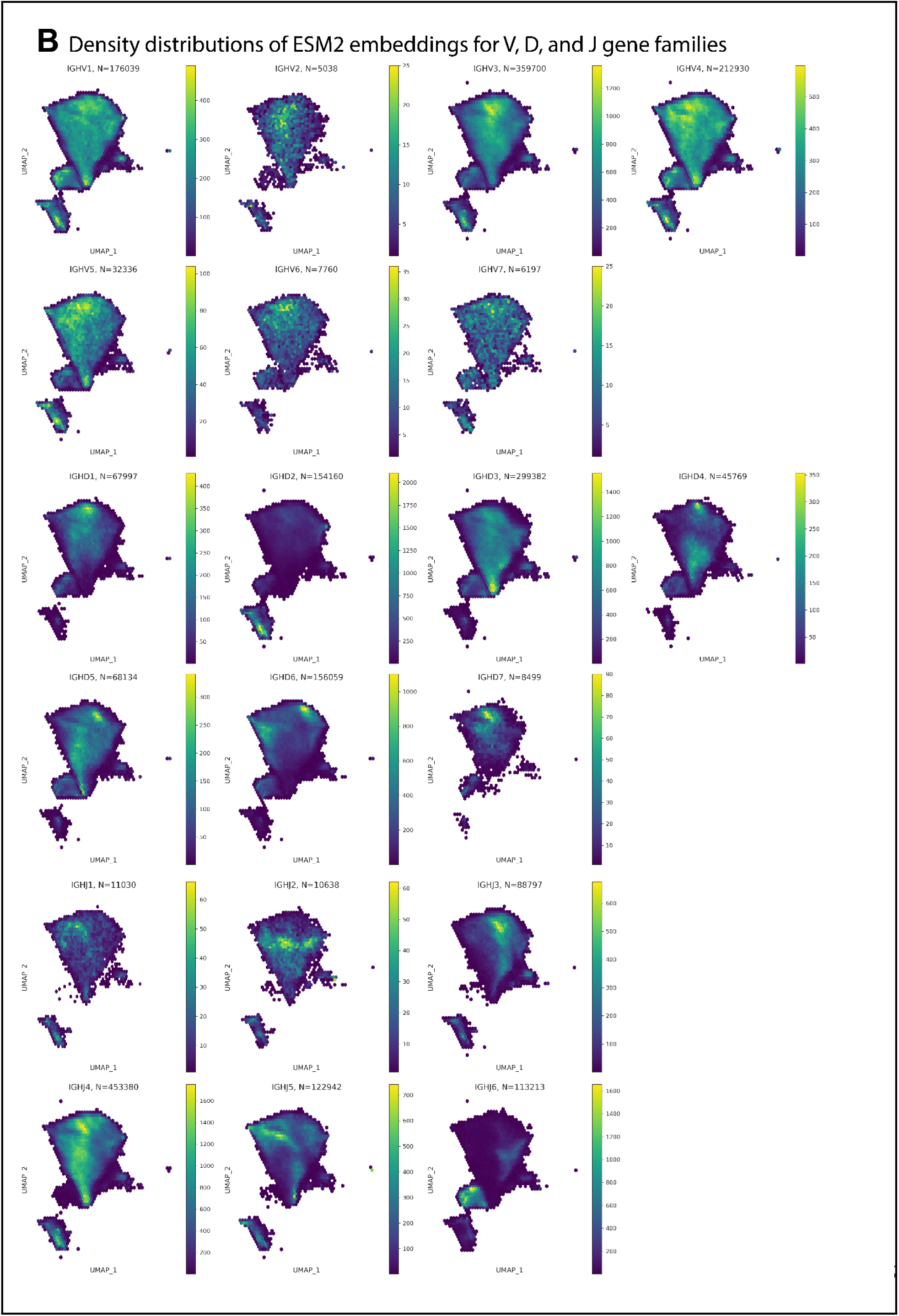
Residual V(D)J gene bias persists in PLM embeddings of HCDR3 sequences. The plots consist of hexbin density maps of A) AB2 and B) ESM2 embeddings from the iReceptor dataset. Separate subplots show distributions for sequences grouped by the V, D, or J gene family. Hexbin coloring reflects local point density. Clustering by J and D families is clearly visible, with weaker but detectable separation for V families. These patterns indicate that PLM embeddings retain residual signatures of germline gene usage, introducing a bias that may confound repertoire-level comparisons. **Refers to main figure:** Fig. 2

**Supplementary figure 14:**
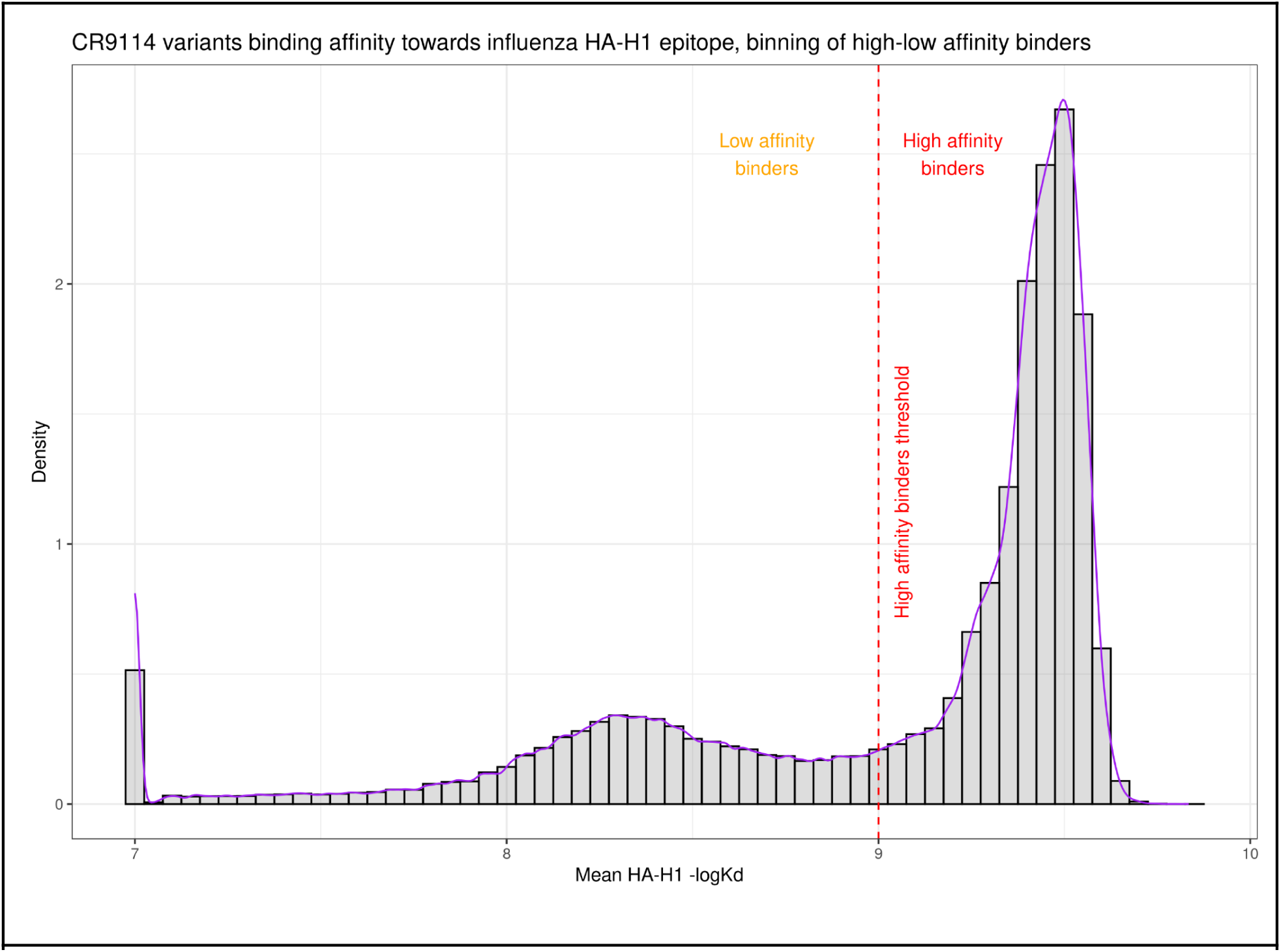
Distribution of CR9114 variant binding affinities against influenza HA-H1 DMS dataset. Histogram representing the CR9114 antibody variants against influenza HA-H1 epitope^78^ from the influenza HA-Shanker dataset used in Fig. 4. As no clear class separation was defined by the authors, we set a threshold at –log(Kd) = 9 to distinguish high from low affinity binders. **Refers to main Figure:** Fig. 4

**Supplementary figure 15:**
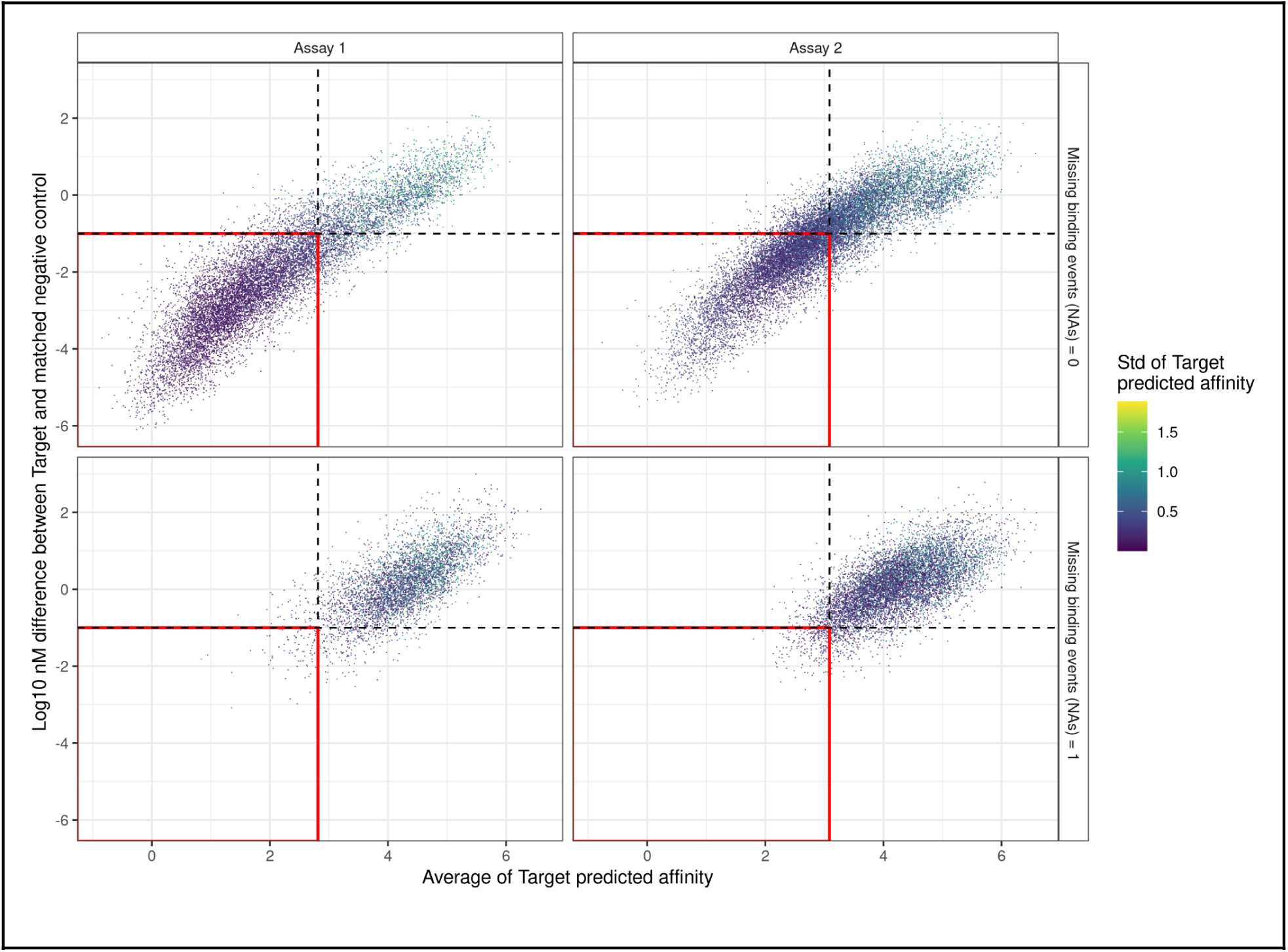
High/low affinity binder binning strategy in the Alphaseq HR2 SARS-CoV-2 dataset. Scatterplots show the gating strategy used to separate antibodies into high and low-binder classes. The *y*-axis represents the difference in binding affinity (log10 Kd, nM) between each antibody’s mean affinity toward the target and the mean affinity of matched negative controls. The *x*-axis shows the average affinity of each antibody toward the target. The red square highlights the region defined by (i) a target–control difference below –1 and (ii) an average binding affinity below the 5th quantile of negative-control affinity scores. The column facets represent two non-overlapping Alphaseq assays that generated the datapoints. Since the two assays exhibited different levels of technical variability, results are shown separately. Faceting rows indicate the number of missing binding events for each antibody variant. A detailed description of the gating strategy is provided in the Methods section. **Refers to main Figure:** Fig. 4

## References

1. Chi, H., Pepper, M. & Thomas, P. G. Principles and therapeutic applications of adaptive immunity. Cell 187, 2052–2078 (2024).

2. Kovaltsuk, A. et al. Observed Antibody Space: A Resource for Data Mining Next-Generation Sequencing of Antibody Repertoires. J. Immunol. 201, 2502–2509 (2018).

3. Corrie, B. D. et al. iReceptor: A platform for querying and analyzing antibody/B-cell and T-cell receptor repertoire data across federated repositories. Immunol. Rev. 284, 24–41 (2018).

4. Omer, A. et al. VDJbase: an adaptive immune receptor genotype and haplotype database. Nucleic Acids Res. 48, D1051–D1056 (2020).

5. Tonegawa, S. Somatic generation of antibody diversity. Nature 302, 575–581 (1983).

6. Georgiou, G. et al. The promise and challenge of high-throughput sequencing of the antibody repertoire. Nat. Biotechnol. 32, 158–168 (2/2014).

7. Mhanna, V. et al. Adaptive immune receptor repertoire analysis. Nature Reviews Methods Primers 4, 1–25 (2024).

8. Talmage, D. W. Immunological specificity, unique combinations of selected natural globulins provide an alternative to the classical concept: Unique combinations of selected natural globulins provide an alternative to the classical concept. Science 129, 1643–1648 (1959).

9. Wucherpfennig, K. W. et al. Polyspecificity of T cell and B cell receptor recognition. Semin. Immunol. 19, 216–224 (2007).

10. Weber, C. R. et al. Reference-based comparison of adaptive immune receptor repertoires. Cell Rep Methods 2, 100269 (2022).

11. Ostrovsky-Berman, M., Frankel, B., Polak, P. & Yaari, G. Immune2vec: Embedding B/T Cell Receptor Sequences in ℝ Using Natural Language Processing. Front Immunol 12, 680687 (2021).

12. Ünlü, A., Ulusoy, E., Yiğit, M. G., Darcan, M. & Doğan, T. Protein language models for predicting drug-target interactions: Novel approaches, emerging methods, and future directions. Curr. Opin. Struct. Biol. 91, 103017 (2025).

13. Wang, M., Patsenker, J., Li, H., Kluger, Y. & Kleinstein, S. H. Language model-based B cell receptor sequence embeddings can effectively encode receptor specificity. Nucleic Acids Res. 52, 548–557 (2024).

14. Pantolini, L. et al. Embedding-based alignment: combining protein language models with dynamic programming alignment to detect structural similarities in the twilight-zone. Bioinformatics 40, (2024).

15. Yeung, W. et al. Tree visualizations of protein sequence embedding space enable improved functional clustering of diverse protein superfamilies. Brief. Bioinform. 24, bbac619 (2023).

16. Dickson, A. & Mofrad, M. R. K. Fine-tuning protein embeddings for functional similarity evaluation. Bioinformatics 40, btae445 (2024).

17. Gonzales, M. E. M., Ureta, J. C. & Shrestha, A. M. S. Protein embeddings improve phage-host interaction prediction. PLoS One 18, e0289030 (2023).

18. Johnson, S. R., Peshwa, M. & Sun, Z. Sensitive remote homology search by local alignment of small positional embeddings from protein language models. Elife 12, (2024).

19. Wang, M., Patsenker, J., Li, H., Kluger, Y. & Kleinstein, S. H. Supervised fine-tuning of pre-trained antibody language models improves antigen specificity prediction. PLoS Comput. Biol. 21, e1012153 (2025).

20. Rao, R., et al. Evaluating protein transfer learning with TAPE. arXiv [cs.LG] (2019) doi:10.1101/676825.

21. Elnaggar, A., et al. ProtTrans: Towards cracking the language of life’s code through self-supervised deep learning and high performance computing. arXiv [cs.LG] (2020).

22. Villegas-Morcillo, A., Gomez, A. M. & Sanchez, V. An analysis of protein language model embeddings for fold prediction. Brief. Bioinform. 23, bbac142 (2022).

23. Matsen, F. A., 4th et al. A sitewise model of natural selection on individual antibodies via a transformer-encoder. Mol. Biol. Evol. 42, msaf186 (2025).

24. Richardson, E., Willemsen, L., Shinde, P., Nielsen, M. & Peters, B. Is the vaccination-induced B cell receptor repertoire predictable? Immunoinformatics (Amst.) 100057 (2025).

25. Bashour, H. et al. Biophysical cartography of the native and human-engineered antibody landscapes quantifies the plasticity of antibody developability. *Commun*. Biol. 7, 922 (2024).

26. Vu, M. H. et al. Linguistically inspired roadmap for building biologically reliable protein language models. Nat Mach Intell 5, 485–496 (2023).

27. Vu, M. H. et al. Linguistics-based formalization of the antibody language as a basis for antibody language models. Nat. Comput. Sci. 4, 412–422 (2024).

28. Simon, E. & Zou, J. InterPLM: discovering interpretable features in protein language models via sparse autoencoders. Nat. Methods 22, 2107–2117 (2025).

29. Johnson, M. M. et al. Nucleotide context models outperform protein language models for predicting antibody affinity maturation. PLoS Comput. Biol. 21, e1013758 (2025).

30. Dounas, A., Cotet, T.-S. & Yermanos, A. Learning immune receptor representations with protein language models. arXiv [q-bio.QM] (2024).

31. Pertseva, M., Follonier, O., Scarcella, D. & Reddy, S. T. TCR clustering by contrastive learning on antigen specificity. Brief. Bioinform. 25, (2024).

32. Kenlay, H. et al. Large scale paired antibody language models. PLoS Comput. Biol. 20, e1012646 (2024).

33. Leem, J., Mitchell, L. S., Farmery, J. H. R., Barton, J. & Galson, J. D. Deciphering the language of antibodies using self-supervised learning. Patterns 3, 100513 (2022).

34. Burbach, S. M. & Briney, B. Improving antibody language models with native pairing. arXiv [q-bio.BM] (2023).

35. Jing, H. et al. Accurate prediction of antibody function and structure using bio-inspired antibody language model. Brief. Bioinform. 25, bbae245 (2024).

36. Wu, K., et al. TCR-BERT: learning the grammar of T-cell receptors for flexible antigen-xbinding analyses. bioRxiv (2021) doi:10.1101/2021.11.18.469186.

37. Olsen, T. H., Moal, I. H. & Deane, C. M. AbLang: an antibody language model for completing antibody sequences. Bioinform Adv 2, vbac046 (2022).

38. Ruffolo, J. A., Gray, J. J. & Sulam, J. Deciphering antibody affinity maturation with language models and weakly supervised learning. arXiv [q-bio.BM] (2021).

39. Singh, R. et al. Learning the language of antibody hypervariability. Proc. Natl. Acad. Sci. U. S. A. 122, (2025).

40. Marcou, Q., Mora, T. & Walczak, A. M. High-throughput immune repertoire analysis with IGoR. Nat. Commun. 9, 561 (2018).

41. Massoni-Badosa, R. et al. An atlas of cells in the human tonsil. Immunity 57, 379–399.e18 (2024).

42. Reed, A. D. et al. A single-cell atlas enables mapping of homeostatic cellular shifts in the adult human breast. Nat. Genet. 56, 652–662 (2024).

43. Barennes, P. et al. Benchmarking of T cell receptor repertoire profiling methods reveals large systematic biases. Nat. Biotechnol. 39, 236–245 (2021).

44. Lin, Z. et al. Evolutionary-scale prediction of atomic-level protein structure with a language model. Science 379, 1123–1130 (2023).

45. Barton, J., Galson, J. D. & Leem, J. Enhancing antibody language models with structural information. bioRxiv (2024) doi:10.1101/2023.12.12.569610.

46. Raybould, M. I. J. et al. The Observed T Cell Receptor Space database enables paired-chain repertoire mining, coherence analysis, and language modeling. Cell Rep. 43, 114704 (2024).

47. Vita, R. et al. The Immune Epitope Database (IEDB): 2024 update. Nucleic Acids Res. 53, D436–D443 (2025).

48. Raybould, M. I. J., Kovaltsuk, A., Marks, C. & Deane, C. M. CoV-AbDab: the coronavirus antibody database. Bioinformatics 37, 734–735 (2021).

49. Briney, B., Inderbitzin, A., Joyce, C. & Burton, D. R. Commonality despite exceptional diversity in the baseline human antibody repertoire. Nature 566, 393–397 (2019).

50. Chinery, L. et al. Baselining the Buzz Trastuzumab-HER2 Affinity, and Beyond. bioRxiv 2024.03.26.586756 (2024) doi:10.1101/2024.03.26.586756.

51. Porebski, B. T. et al. Rapid discovery of high-affinity antibodies via massively parallel sequencing, ribosome display and affinity screening. *Nat*. Biomed. Eng. 8, 214–232 (2024).

52. Greiff, V., Miho, E., Menzel, U. & Reddy, S. T. Bioinformatic and statistical analysis of adaptive immune repertoires. Trends Immunol. 36, 738–749 (2015).

53. Trepel, F. Number and distribution of lymphocytes in man. A critical analysis. Klin. Wochenschr. 52, 511–515 (1974).

54. Farber, D. L., Yudanin, N. A. & Restifo, N. P. Human memory T cells: generation, compartmentalization and homeostasis. Nat. Rev. Immunol. 14, 24–35 (2014).

55. Preuer, K., Renz, P., Unterthiner, T., Hochreiter, S. & Klambauer, G. Fréchet ChemNet distance: A metric for generative models for molecules in drug discovery. J. Chem. Inf. Model. 58, 1736–1741 (2018).

56. Borji, A. Pros and cons of GAN evaluation measures. Comput. Vis. Image Underst. 179, 41–65 (2019).

57. Maaten, L. & Hinton, G. E. Visualizing Data using t-SNE. Journal of Machine Learning Research 9, 2579–2605 (2008).

58. Sun, Y. et al. A comprehensive survey of dimensionality reduction and clustering methods for single-cell and spatial transcriptomics data. Brief. Funct. Genomics 23, 733–744 (2024).

59. Raybould, M. I. J. et al. The Observed T cell receptor Space database enables paired-chain repertoire mining, coherence analysis and language modelling. Immunology (2024).

60. Hubert, L. & Arabie, P. Comparing partitions. J. Classif. 2, 193–218 (1985).

61. Sethna, Z. et al. Population variability in the generation and selection of T-cell repertoires. PLoS Comput. Biol. 16, e1008394 (2020).

62. Isacchini, G., Walczak, A. M., Mora, T. & Nourmohammad, A. Deep generative selection models of T and B cell receptor repertoires with soNNia. Proc Natl Acad Sci U S A 118, (2021).

63. Hopp, C. S., et al. Atypical B cells up-regulate costimulatory molecules during malaria and secrete antibodies with T follicular helper cell support. Sci. Immunol. 7, eabn1250 (2022).

64. Chen, E. C. et al. Systematic analysis of human antibody response to ebolavirus glycoprotein shows high prevalence of neutralizing public clonotypes. Cell Rep. 42, 112370 (2023).

65. Roy, B. et al. High-throughput single-cell analysis of B cell receptor usage among autoantigen-specific plasma cells in celiac disease. J. Immunol. 199, 782–791 (2017).

66. Chronister, W. D. et al. TCRMatch: Predicting T-cell receptor specificity based on sequence similarity to previously characterized receptors. Front. Immunol. 12, 640725 (2021).

67. Lanzarotti, E., Marcatili, P. & Nielsen, M. T-cell receptor cognate target prediction based on paired α and β chain sequence and structural CDR loop similarities. Front. Immunol. 10, 2080 (2019).

68. Mayer-Blackwell, K. et al. TCR meta-clonotypes for biomarker discovery with tcrdist3 enabled identification of public, HLA-restricted clusters of SARS-CoV-2 TCRs. Elife 10, (2021).

69. Wilamowski, J., et al. InterClone: Store, search and cluster adaptive immune receptor repertoires. bioRxiv 2022.07.31.501809 (2022) doi:10.1101/2022.07.31.501809.

70. Abbate, M. F. et al. Computational detection of antigen-specific B cell receptors following immunization. Proc. Natl. Acad. Sci. U. S. A. 121, e2401058121 (2024).

71. Ye, C., Hu, W. & Gaeta, B. Prediction of antibody-antigen binding via machine learning: Development of data sets and evaluation of methods. JMIR Bioinform. Biotech. 3, e29404 (2022).

72. Mayer, A. & Callan, C. G., Jr. Measures of epitope binding degeneracy from T cell receptor repertoires. Proc. Natl. Acad. Sci. U. S. A. 120, e2213264120 (2023).

73. Dash, P. et al. Quantifiable predictive features define epitope-specific T cell receptor repertoires. Nature 547, 89–93 (2017).

74. Marro, S. et al. Language models are implicitly continuous. arXiv [cs.CL] (2025).

75. Pedregosa, F. et al. Scikit-learn: Machine Learning in Python. J. Mach. Learn. Res.

76. Backurs, A. & Indyk, P. Edit Distance Cannot Be Computed in Strongly Subquadratic Time (unless SETH is false). arXiv [cs.CC] (2014).

77. Heinzinger, M., et al. Bilingual language model for protein sequence and structure. NAR Genom. Bioinform. 6, lqae150 (2024).

78. Shanker, V. R., Bruun, T. U. J., Hie, B. L. & Kim, P. S. Unsupervised evolution of protein and antibody complexes with a structure-informed language model. Science 385, 46–53 (2024).

79. Engelhart, E. et al. A dataset comprised of binding interactions for 104,972 antibodies against a SARS-CoV-2 peptide. Sci. Data 9, 653 (2022).

80. Luo, Z. et al. Interpretable feature extraction and dimensionality reduction in ESM2 for protein localization prediction. Brief. Bioinform. 25, bbad534 (2024).

81. Garbas, L., Ploner, M. & Akbik, A. TransformerRanker: A tool for efficiently finding the best-suited language models for downstream classification tasks. arXiv [cs.CL] (2024).

82. Mason, D. M. et al. Optimization of therapeutic antibodies by predicting antigen specificity from antibody sequence via deep learning. *Nat*. Biomed. Eng. 5, 600–612 (2021).

83. https://papers.nips.cc/paper_files/paper/2017/hash/8a1d694707eb0fefe65871369074926d-Abstract.html.

84. Kilgour, K., Zuluaga, M., Roblek, D. & Sharifi, M. Fr’echet Audio Distance: A metric for evaluating music enhancement algorithms. arXiv [eess.AS] (2018).

85. Unterthiner, T., et al. Towards accurate generative models of video: A new metric & challenges. arXiv [cs.CV] (2018).

86. Olson, B. J., Schattgen, S. A., Thomas, P. G., Bradley, P. & Matsen, F. A., Iv. Comparing T cell receptor repertoires using optimal transport. PLoS Comput. Biol. 18, e1010681 (2022).

87. Rissom, P. F. et al. Decoding protein language models: insights from embedding space analysis. Bioinformatics (2024).

88. Rives, A. et al. Biological structure and function emerge from scaling unsupervised learning to 250 million protein sequences. Proc. Natl. Acad. Sci. U. S. A. 118, e2016239118 (2021).

89. OpenAI et al. GPT-4 Technical Report. arXiv [cs.CL] (2023).

90. Comanici, G. et al. Gemini 2.5: Pushing the frontier with advanced reasoning, multimodality, long context, and next generation agentic capabilities. arXiv [cs.CL] (2025).

91. Introducing Claude 4. https://www.anthropic.com/news/claude-4.

92. Wang, L., et al. A comprehensive review of protein language models. arXiv [q-bio.BM] (2025).

93. Neyestanak, M. S. et al. Data-optimal scaling of paired antibody language models. bioRxivorg 2025.09.02.673765 (2025) doi:10.1101/2025.09.02.673765.

94. Nijkamp, E., Ruffolo, J., Weinstein, E. N., Naik, N. & Madani, A. ProGen2: Exploring the Boundaries of Protein Language Models. (2023).

95. Rissom, P. F. et al. Decoding protein language models: insights from embedding space analysis.bioRxiv (2024) doi:10.1101/2024.06.21.600139.

96. Analysis of antigen-binding proteins. World Patent (2025).

97. Hie, B. L., Yang, K. K. & Kim, P. S. Evolutionary velocity with protein language models predicts evolutionary dynamics of diverse proteins. Cell Syst. 13, 274–285.e6 (2022).

98. Radovanović, M., Nanopoulos, A. & Ivanović, M. Hubs in space: Popular nearest neighbors in high-dimensional data. J. Mach. Learn. Res. 11, 2487–2531 (2010).

99. Skean, O., et al. Layer by layer: Uncovering hidden representations in language models. arXiv [cs.LG] (2025).

100. Lad, V., Lee, J. H., Gurnee, W. & Tegmark, M. The remarkable robustness of LLMs: Stages of inference? arXiv [cs.LG] (2024).

101. Deutschmann, N. et al. Do domain-specific protein language models outperform general models on immunology-related tasks? Immunoinformatics (Amst.) 14, 100036 (2024).

102. Kobak, D. & Berens, P. The art of using t-SNE for single-cell transcriptomics. Nat. Commun. 10, 5416 (2019).

103. Becht, E. et al. Dimensionality reduction for visualizing single-cell data using UMAP. Nat. Biotechnol. 37, 38–44 (2018).

104. Glänzer, W. S., Reddy, S. T. & Yermanos, A. Revealing bias in antibody language models through systematic training data processing with OAS-explore. in NeurIPS 2025 2nd Workshop on Multi-modal Foundation Models and Large Language Models for Life Sciences (2025).

105. Talaei, M. et al. CDR-aware masked language models for paired antibodies enable state-of-the-art binding prediction. bioRxiv 2025.10.31.685149 (2025) doi:10.1101/2025.10.31.685149.

106. Gujral, O., Bafna, M., Alm, E. & Berger, B. Sparse autoencoders uncover biologically interpretable features in protein language model representations. Proc. Natl. Acad. Sci. U. S. A. 122, e2506316122 (2025).

107. Schütze, K., Heinzinger, M., Steinegger, M. & Rost, B. Nearest neighbor search on embeddings rapidly identifies distant protein relations. Front. Bioinform. 2, 1033775 (2022).

108. Steck, H., Ekanadham, C. & Kallus, N. Is cosine-similarity of embeddings really about similarity? arXiv [cs.IR] (2024) doi:10.1145/3589335.3651526.

109. Peng, D., Gui, Z. & Wu, H. Interpreting the curse of dimensionality from distance concentration and manifold effect. arXiv [cs.LG] (2023).

110. Canessa, E., Chaigneau, S. E., Moreno, S. & Lagos, R. Informational content of cosine and other similarities calculated from high-dimensional Conceptual Property Norm data. Cogn. Process. 21, 601–614 (2020).

111. Iovino, B. G. & Ye, Y. Protein embedding based alignment. BMC Bioinformatics 25, 85 (2024).

112. Kulikova, A. V., Parker, J. K., Davies, B. W. & Wilke, C. O. Semantic search using protein large language models detects class II microcins in bacterial genomes. mSystems 9, e0104424 (2024).

113. Odrzywolek, K. et al. Deep embeddings to comprehend and visualize microbiome protein space. Sci. Rep. 12, 10332 (2022).

114. O’Donnell, T. J. et al. Reading the repertoire: Progress in adaptive immune receptor analysis using machine learning. Cell Syst. 15, 1168–1189 (2024).

115. Ursu, E. et al. Training data composition determines machine learning generalization and biological rule discovery. *Nat*. Mach. Intell. 7, 1206–1219 (2025).

116. Ye, J., Ma, N., Madden, T. L. & Ostell, J. M. IgBLAST: an immunoglobulin variable domain sequence analysis tool. Nucleic Acids Res. 41, W34–40 (2013).

117. Pavlović, M. et al. The immuneML ecosystem for machine learning analysis of adaptive immune receptor repertoires. *Nat*. Mach. Intell. 3, 936–944 (2021).

118. Su, J. et al. RoFormer: Enhanced transformer with Rotary Position Embedding. arXiv [cs.CL] (2021).

119. Suzek, B. E. et al. UniRef clusters: a comprehensive and scalable alternative for improving sequence similarity searches. Bioinformatics 31, 926–932 (2015).

120. Van Der Maaten, L. & Hinton, G. Visualizing data using t-SNE. Journal of Machine Learning Research 9, 2579–2605 (2008).

121. Poličar, P. G., Stražar, M. & Zupan, B. openTSNE: a modular Python library for t-SNE dimensionality reduction and embedding. bioRxiv (2019) doi:10.1101/731877.

122. Spisak, N., Athènes, G., Dupic, T., Mora, T. & Walczak, A. M. Combining mutation and recombination statistics to infer clonal families in antibody repertoires. Elife 13, e86181 (2024).

123. Ripley, B. & Venables, B. MASS: Support functions and datasets for Venables and ripley’s MASS. CRAN: Contributed Packages The R Foundation 10.32614/cran.package.mass (2009).

124. Zhong, J., niccolocard & Habib-Bashour. Csi-Greifflab/pepe-Cli: Release v1.0.5. (Zenodo, 2025). doi:10.5281/ZENODO.16273335.

