## Supplementary Table 1 for "Quantitative mapping of antigen specificity in adaptive immune repertoire embedding spaces": Table_S1.pdf

| Dataset name in the paper | Total number of sequences (original source) | Total number of sequences used in our study | Topic | Organisms | BCR/TCR | Chains | Specificity labels | Reference |
|---|---|---|---|---|---|---|---|---|
| | Number of sequences | Number of sequences | Eg. Disease/Vaccine, mixed... | Eg. mouse, human... | BCR / TCR | Eg. paired / heavy / HCDR3... | Eg. Yes/No | |
| iReceptor | 616,095,579 | 2,151,099 | Mixed | human | BCR | fullchain heavy + HCDR3 | No | https://gateway.ireceptor.org |
| Wang | 794,620 | 794,620 | Mixed | human | BCR | HCDR3 | No | https://academic.oup.com/nar/article/52/2/548/7477518 |
| HER2 Chinery | 349,893 | 344,688 | Variants of HER2 binder | synthetic- derived from hum | BCR | paired (identical light chains) | Yes | https://www.biorxiv.org/content/10.1101/2024.03.26.586756v1 |
| Sars-Cov-2 RBD CovAb-Dab | 12,535 | 7,235 | SARS-CoV-2 | human | BCR | paired | Yes (Antigen; Epitope) | https://opig.stats.ox.ac.uk/webapps/covabdab/ |
| Influenza | 181 | 171 | Influenza from IEDB | human | BCR | paired | Yes | https://iedb.org |
| TG2+ | 1,454 | 1,454 | Celiac disease | human | BCR | heavy | Yes | https://academic.oup.com/jimmunol/article-abstract/199/2/782/796 |
| Sars-Cov-2 HR2 Engelhart | 104,972 | 65,619 | SARS-CoV-2 | synthetic | BCR | paired | No (binding affinity) | https://www.nature.com/articles/s41597-022-01779-4 |
| Briney | 364,000,000 | 18,000,000 | Healthy donors | human | BCR | heavy + light + V/D/J/ gene | yes | https://www.nature.com/articles/s41586-019-0879-y/ |
| Ebola | 10,326 | 10,326 | Ebola | human | BCR | V_gene+J_gene+H-CDR3+L-CDR3+ | Yes | https://doi.org/10.1016/j.celrep.2023.112370 |
| Influenza HA-Shanker | 69,676 | 69,676 | deep mutational scanning - influenza | synthetic | BCR | paired | No (binding affinity) | https://www.science.org/doi/10.1126/science.adk8946 |
| OTS | 5,913,192 | 399,990 | Mixed | human | TCR | paired alpha/beta and gamma/delta | No | https://opig.stats.ox.ac.uk/webapps/ots/ |
| HER2 Porebski | 24,970 | 24,970 | LLM-generated HER2-binders | synthetic | BCR | HCDR3 | No (binding affinity) | https://www.nature.com/articles/s41551-023-01093-3 |
| HIV | 284 | 284 | HIV protein-specific antibodies | human | BCR | paired | yes | https://iedb.org |
| Postselection | 1,000,000 | 1,000,000 | simulated heavy chain dataset | simulated (in silico) | BCR | V_gene+J_gene+HCDR3 | No | https://github.com/statbiophys/soNNia |
| Malaria | 1,286 | 1,286 | Pf specific b cells | human | BCR | HCDR3 | yes | https://pmc.ncbi.nlm.nih.gov/articles/PMC11132112/ |
| Influenza | 1,076 | 1,076 | HA specific b cells | human | BCR | HCDR3 | yes | https://pmc.ncbi.nlm.nih.gov/articles/PMC11132112/ |
