## Supplementary Table 2 for "Quantitative mapping of antigen specificity in adaptive immune repertoire embedding spaces": Table_S2.pdf

| Figure panel | Model | Downsample_size | Dataset | Sequence context window | Effect sizes | | | | | |
|---|---|---|---|---|---|---|---|---|---|---|
| | | | | | Pooled_vs_Attention | Pooled_vs_Unpooled | Attention_vs_Unpooled | LD_vs_Pooled | LD_vs_Attention | LD_vs_Unpooled |
| 4C | Antiberta2-cssp | - | HER2 Chinery | all_cdrh | 0.0583 | 0.2321 | 0.1773 | 0.2844 | 0.2215 | 0.0302 |
| 4C | ESM2 | - | HER2 Chinery | all_cdrh | 0.0373 | 0.2580 | 0.2920 | 0.2881 | 0.3227 | 0.0557 |
| 4C | Antiberta2-cssp | - | HER2 Chinery | hcdr3_extracted | 0.0887 | 0.2598 | 0.3415 | 0.2605 | 0.3913 | 0.1204 |
| 4C | ESM2 | - | HER2 Chinery | hcdr3_extracted | 0.0154 | 0.2704 | 0.2890 | 0.2880 | 0.3092 | 0.0800 |
| 4C | Antiberta2-cssp | - | HER2 Chinery | hcdr3_only | 0.0435 | 0.2199 | 0.2702 | 0.2759 | 0.3414 | 0.0240 |
| 4C | ESM2 | - | HER2 Chinery | hcdr3_only | 0.0488 | 0.3130 | 0.2443 | 0.3719 | 0.2997 | 0.0007 |
| 4C | Antiberta2-cssp | - | HER2 Chinery | heavy_chain | 0.1701 | 0.2881 | 0.1085 | 0.3262 | 0.0848 | 0.0691 |
| 4C | ESM2 | - | HER2 Chinery | heavy_chain | 0.0741 | 0.2350 | 0.1515 | 0.3004 | 0.2142 | 0.0031 |
| 4C | Antiberta2-cssp | - | HER2 Chinery | paired_chain | 0.1391 | 0.2364 | 0.0894 | 0.2492 | 0.0690 | 0.0920 |
| 4C | ESM2 | - | HER2 Chinery | paired_chain | 0.1136 | 0.2734 | 0.1470 | 0.3367 | 0.2083 | 0.0035 |
| 4D | Antiberta2-cssp | 25000 | HER2 Chinery | hcdr3_only | -0.0956 | -0.1651 | -0.2398 | 0.3331 | -0.3946 | 0.1237 |
| 4D | ESM2 | 25000 | HER2 Chinery | hcdr3_only | -0.0014 | -0.3244 | -0.3031 | 0.4770 | -0.4523 | 0.1048 |
| 4D | Antiberta2-cssp | 10000 | HER2 Chinery | hcdr3_only | -0.0899 | -0.1642 | -0.2354 | 0.2814 | -0.3395 | 0.0980 |
| 4D | ESM2 | 10000 | HER2 Chinery | hcdr3_only | -0.0310 | -0.3028 | -0.3108 | 0.3968 | -0.3998 | 0.0705 |
| 4D | Antiberta2-cssp | 5000 | HER2 Chinery | hcdr3_only | -0.1403 | -0.1276 | -0.2445 | 0.2093 | -0.3204 | 0.0668 |
| 4D | ESM2 | 5000 | HER2 Chinery | hcdr3_only | -0.0778 | -0.2824 | -0.3459 | 0.3732 | -0.4342 | 0.0808 |
| 4D | Antiberta2-cssp | 2500 | HER2 Chinery | hcdr3_only | -0.2403 | -0.1382 | -0.3545 | 0.3273 | -0.5068 | 0.1847 |
| 4D | ESM2 | 2500 | HER2 Chinery | hcdr3_only | -0.0983 | -0.2032 | -0.3150 | 0.4063 | -0.5259 | 0.1949 |
| 4E | Antiberta2-cssp | - | HER2 Porebski | hcdr3_only | 0.2569 | 0.1686 | 0.0874 | 2.0598 | 1.8604 | 1.9118 |
| 4E | ESM2 | - | HER2 Porebski | hcdr3_only | 0.1462 | 0.2362 | 0.0724 | 3.3839 | 2.6824 | 2.9725 |
| 4E | Antiberta2-cssp | - | Influenza HA | heavy_chain | 0.5724 | 0.0721 | 0.6576 | 0.6318 | 1.0182 | 0.6544 |
| 4E | ESM2 | - | Influenza HA | heavy_chain | 0.2835 | 0.0288 | 0.6714 | 1.5342 | 3.2575 | 2.2166 |
| 4E | Antiberta2-cssp | - | SARS-CoV-2 HR2 | paired_chain | 0.2907 | 0.3142 | 0.0072 | 0.6645 | 0.4068 | 0.4098 |
| 4E | ESM2 | - | SARS-CoV-2 HR2 | paired_chain | 0.1210 | 0.1870 | 0.0529 | 0.3921 | 0.2789 | 0.2523 |
| 4E | Antiberta2-cssp | - | SARS-CoV-2 RBD | paired_chain | 0.3196 | 0.3344 | 0.0328 | 1.5882 | 3.0710 | 2.8104 |
| 4E | ESM2 | - | SARS-CoV-2 RBD | paired_chain | 0.3612 | 0.2079 | 0.2388 | 1.6307 | 3.8875 | 2.6940 |
